# Online self-evaluation of fMRI-based neurofeedback performance

**DOI:** 10.1101/2021.08.20.457108

**Authors:** Santiago Muñoz-Moldes, Anita Tursic, Michael Lührs, Judith Eck, Amaia Benitez Andonegui, Judith Peters, Axel Cleeremans, Rainer Goebel

**Affiliations:** Consciousness, Cognition and Computation group, Center for Research in Cognition & Neuroscience, Faculty of Psychology and Education, Université Libre de Bruxelles, Brussels, Belgium; Department of Psychology, University of Cambridge, Cambridge, United Kingdom; Brain Innovation B.V., Research Department, Maastricht, the Netherlands; Department of Cognitive Neuroscience, Faculty of Psychology and Neuroscience, Maastricht University, Maastricht, the Netherlands; Netherlands Institute for Neuroscience, Amsterdam, the Netherlands

**Keywords:** neurofeedback, functional magnetic resonance imaging (fMRI), self-regulation, metacognition, confidence, motor imagery

## Abstract

This study explores the subjective evaluation of supplementary motor area (SMA) regulation performance in a real-time functional magnetic resonance imaging neurofeedback (fMRI-NF) task. In fMRI-NF, people learn how to self-regulate their brain activity by performing mental actions to achieve a certain target level of blood-oxygen-level-dependent (BOLD) activation. This setup offers the possibility to study performance monitoring in the absence of somatosensory feedback. Here, we studied two types of self-evaluation expressed before receiving neurofeedback: performance predictions and perceived confidence in the prediction judgement. We hypothesized that throughout learning, participants would (1) improve the precision of their performance predictions about the actual changes in their BOLD response, and (2) that reported confidence would progressively increase with improved metacognitive precision. Participants completed three sessions of SMA regulation in a 7T fMRI scanner, performing a drawing motor imagery task. During each trial, they modulated their mental drawing strategy to achieve one of two different levels of target fMRI signal change. They then reported a performance prediction and their confidence in the prediction before receiving delayed BOLD-activation feedback. Results show that participants’ performance predictions improved with learning throughout the three sessions, and that these improvements were not driven exclusively by their knowledge of previous performance. Confidence reports on the other hand showed no change throughout training and did not differentiate between the better and worse predictions. In addition to shedding light on mechanisms of internal monitoring during neurofeedback training, these results also point to a dissociation between self-evaluation of performance and corresponding reported confidence in the presence of feedback.

## 1 Introduction

Neurofeedback is a special type of biofeedback that enables self-regulation of one’s brain activity and can be used either by healthy participants aiming to improve their cognitive performance, or as an intervention strategy for symptom improvement in clinical populations (for a review see Sitaram et al., 2017). Neurofeedback signals based on real-time functional magnetic resonance imaging (fMRI-NF) offer high regional specificity (Sulzer et al., 2013; Weiskopf et al., 2004, 2003) and can in turn achieve higher precision in behavioral outcomes compared to other modalities, such as electroencephalography (EEG) or functional near-infrared spectroscopy (fNIRS) (Scharnowski et al., 2015, 2012; Weiskopf et al., 2004). Using high field (7T) MRI can additionally help to achieve better signal-to-noise ratio in comparison to using more standard, lower MRI field strengths (Torrisi et al., 2018). This can be particularly beneficial during gradual, level-specific self-regulation of the BOLD signal (Sorger et al., 2018; Krause et al., 2017; Sousa et al., 2016), by providing more degrees of freedom for learning self-regulation in the context of neurofeedback and brain-computer interfaces (BCI) than more standard up- or down-regulation.

Although neurofeedback has been shown to successfully help brain activity regulation (for reviews see Thibault et al., 2018; Tursic et al., 2020), the underlying learning mechanisms of neurofeedback remain unclear. It has been hypothesized that they depend on the neurofeedback procedure and on the exact instructions given to the participants, such as the suggestion of explicit regulatory strategies, or the knowledge that feedback depends on one’s brain activation (Muñoz-Moldes & Cleeremans, 2020; Sepulveda et al., 2016; Sitaram et al., 2017; Sulzer et al., 2013). One theory proposes that neurofeedback learning is a form of instrumental conditioning by which a neural response becomes more likely when associated with a reward (Fetz, 2007; Kamiya, 1962; Shibata et al., 2019). Another proposal suggests that feedback exposure allows control of physiological activity by increasing our awareness of internal –otherwise inaccessible–states (Brener, 1977; Brown, 1971; Frederick et al., 2016; Neumann et al., 2003).

The relation between awareness and control not only has important implications for regulation but also for the general applicability, or transfer success, of neurofeedback: indeed, researchers often evaluate performance in transfer trials (i.e., doing the self-regulation task without feedback) as an indicator of how well the learned self-regulation might translate to the real world (Sitaram et al., 2017). Presumably, while regulating without any feedback, participants compute an internal estimate of their ongoing performance (which might then be accompanied by a feeling of confidence) and use this estimate to fine-tune their regulation task, making performance monitoring an important component of successful self-regulation. Several studies indeed reported successful regulation during transfer runs (Young et al., 2017; Orlov et al., 2018; Alegria et al., 2017), including regulation to different levels (Sousa et al., 2016), but none investigated how self-monitoring contributed to this success. Understanding self-evaluation accuracy, and the possible manipulation of such self-evaluations might result in incentivizing learning and in the improvement of clinical applications. It would also, importantly, be useful for our theoretical understanding of how such internal monitoring and associated feelings of confidence are computed.

One crucial component of neurofeedback self-regulation is metacognition. Metacognition – cognition about cognition – can be defined as the self-evaluation of the quality of neuronal evidence (Fleming et al., 2012a; Yeung and Summerfield, 2012). It is associated with performance monitoring and error awareness, two different, but interdependent processes (Metcalfe and Greene, 2007; Miele et al., 2011). In the case of neurofeedback and BCI, metacognitive decisions can be understood as a form of internal monitoring detached from somatosensory feedback, a process that is thus different from the monitoring of executed movements (Schurger et al., 2017). In neurofeedback tasks, two types of self-evaluation can be distinguished: the evaluation of a mental action or signal (i.e., the evaluation of performance or an assessment of the predicted consequence, such as the prediction of the feedback value), and the evaluation of one’s own evaluation (i.e., an assessment of one’s own monitoring performance, such as the confidence of having made an accurate prediction of the feedback). As neurofeedback often requires practice over multiple sessions (Ahn and Jun, 2015), it can be considered a suitable candidate for studying changes in performance prediction and confidence over the course of neurofeedback learning.

The neural architecture supporting metacognition and performance monitoring is still debated, and in particular the extent to which it consists of domain-specific modules (e.g., for perception and memory), of a domain-general component, or both (Faivre et al., 2018; Rouault et al., 2018). Metacognition in perceptual and memory tasks, which are perhaps the closest to neurofeedback, typically involves a frontoparietal network, including posterior medial, ventromedial, and dorsolateral prefrontal cortex (pMPFC, vMPFC, DLPFC, respectively), precuneus and insula (Vaccaro and Fleming, 2018). Previous studies have also related error awareness to activity in posterior medial prefrontal cortex (pMPFC), SMA and insula (Bonini et al., 2014; Ullsperger et al., 2010).

In the present work, we study how performance self-evaluation for self-generated mental actions and its associated confidence evolve during neurofeedback-guided motor imagery training. Participants were trained to adjust their motor imagery to two different intensity levels, meaning that they were asked to reach a certain percentage of their maximal regional activity, rather than to maximize the activity, which is more commonly done in neurofeedback research. The levels were individually defined as a function of their maximum performance in the initial localizer task. They then expressed interleaved performance predictions (i.e., predictions of feedback) and associated judgments of confidence, before receiving intermittent neurofeedback from the SMA region using a 7 Tesla MRI. We hypothesized that as self-regulation performance improved, participants would increase their accuracy in their self-evaluation, as evidenced both by better performance predictions, and a higher match between confidence and prediction accuracy.

## 2 Methods

### 2.1 Participants

Eleven participants were recruited at Maastricht University (Maastricht, the Netherlands) to undergo five training sessions, one per day, and all completed within at most eleven days. To maximize learning and hence the associated potential change in performance prediction and confidence, we recruited participants without previous neurofeedback experience. One participant performed an alternative version of the task with other experimental parameters (P01), another did not finish all sessions (P03), and a third failed to follow the instructions of the task (P10); we thus excluded them from further analysis. The final sample consisted of eight healthy volunteers (four females), aged 25–32 years (*M* = 27.5, *SD* = 2.5), all right-handed, with normal or corrected-to-normal vision and without any history of previous psychiatric or neurological disorders. Note that the original participant labels (P01 – P11) were kept for consistency. Participants provided informed consent and received financial compensation for taking part in the study.

### 2.2 General procedure

The experimental procedure was approved by the *Ethics Review Committee Psychology and Neuroscience* at Maastricht University and was in conformity to the standards of the Declaration of Helsinki (6th revision, 2008). The study consisted of five sessions, during which participants completed trials of a motor imagery task interleaved with self-reports of performance prediction and confidence (see 3.2 for details). The first and fifth session took place with a functional near-infrared spectroscopy (fNIRS) measure outside the scanner and without neurofeedback. The other three sessions (second, third and fourth) were performed in the 7 Tesla fMRI scanner and included intermittent neurofeedback after every trial. Here, we only present data of the neurofeedback fMRI sessions (second to fourth session), hereafter referred to as sessions 1st, 2nd and 3rd.

Each fMRI session lasted approximately two hours and consisted of one anatomical measurement and seven functional runs. The first functional run was an eight-minute localizer scan used to define the target region and the parameters for the neurofeedback runs. The subsequent neurofeedback runs lasted 9 minutes each and included 60 trials of self-regulation in each session (10 trials per run). Hence, each participant performed a total of 180 trials (90 per condition) across the three sessions.

### 2.3 Tasks

#### 2.3.1 Functional localizer task

At the beginning of each session, participants performed a motor imagery and a finger tapping task to define the target region and the thresholds for the neurofeedback task (see 2.5.2 Target region selection). The localizer run consisted of alternating 16-second blocks of mental drawing, finger tapping, and rest. In mental drawing blocks, participants performed mental drawing (see *Instructions for motor imagery in* 2.3). In finger tapping blocks, participants tapped to an auditory signal at 2 Hz with their right index and middle fingers using a button box. The auditory cues were used to keep the display uniform across different tasks and to ensure tapping consistency across sessions and participants. Participants completed eight blocks of drawing, eight blocks of tapping and 17 blocks of rest.

##### Instructions for motor imagery

Participants were informed about the principles of neurofeedback and were given suggestions for the cognitive strategies they could use. We explained that their goal in the experiment was to “learn how to achieve different levels of brain activation by modulating a mental drawing task”. They were suggested to focus on the feelings and sensations associated with the preparation or execution of a movement with their right upper limb, while keeping their eyes open and refraining from overt hand movements (which was visually monitored by an experimenter during the first fNIRS session). It was suggested that the intensity could be modulated by adjusting one aspect of the mental drawing strategy, such as the drawing detail, the speed, the (imagined) pressure exerted or the number of muscles involved (e.g., fingers, wrist, arm, shoulder). Participants were asked to maintain the strategy within a single trial but were otherwise allowed to try different strategies based on their feedback and performance. For the functional localizer tasks, the participants were asked to perform mental drawing at the highest intensity that they could imagine.

#### 2.3.2 Motor imagery neurofeedback task

Stimuli were presented using the Expyriment package (version 0.9.0) for Python (version 2.7.10) (Krause and Lindemann, 2014). Each trial started with a red cross signaling the rest period and the spoken word “rest”. After 16 seconds, the motor imagery period started with a change in color of the cross (to yellow or green) and a simultaneous auditory cue (“six” or “nine”), indicating the target level to be achieved through motor imagery (60% or 90% of the maximum percent-signal change (MaxPSC) defined based on the functional localizer, respectively). The auditory cues were used to keep the display uniform across the two conditions and at the same time keep the memory load associated with the cross color minimal. The indicated target levels were pseudorandomized so that half of the trials in the run requested the participants to regulate to level 60% and half to level 90%. After 16 seconds of continuous motor imagery, an auditory cue “stop” indicated the end of the motor imagery trial. This was followed by a jittered blank screen of 1, 2- or 3-seconds duration (average of 2s). Participants were then shown a horizontal rating scale showing values from 0 to 12, with 6 and 9 representing the two target levels (60% and 90%, respectively). They were asked to report their performance prediction by moving left or right on the scale with two buttons of the button box. After another jittered blank screen (1, 2 or 3 seconds, average of 2), a second scale was presented, and participants were asked to report their confidence in their prediction. The horizontal scale contained values ranging from 50% to 100% increasing in steps of 5%, and two labels, one on the far-left indicating ‘Guess’ and one on the far-right indicating ‘Totally sure’. For both scales, participants had six seconds to respond, and the start position of the cursor was jittered around the midpoint (-4; +4) to prevent motor preparation. Participants were then shown the neurofeedback value on the same 13-point scale for 2 seconds, with an arrow pointing from the target level to the achieved value regulation (see Fig. 1D).

**Figure 1.**
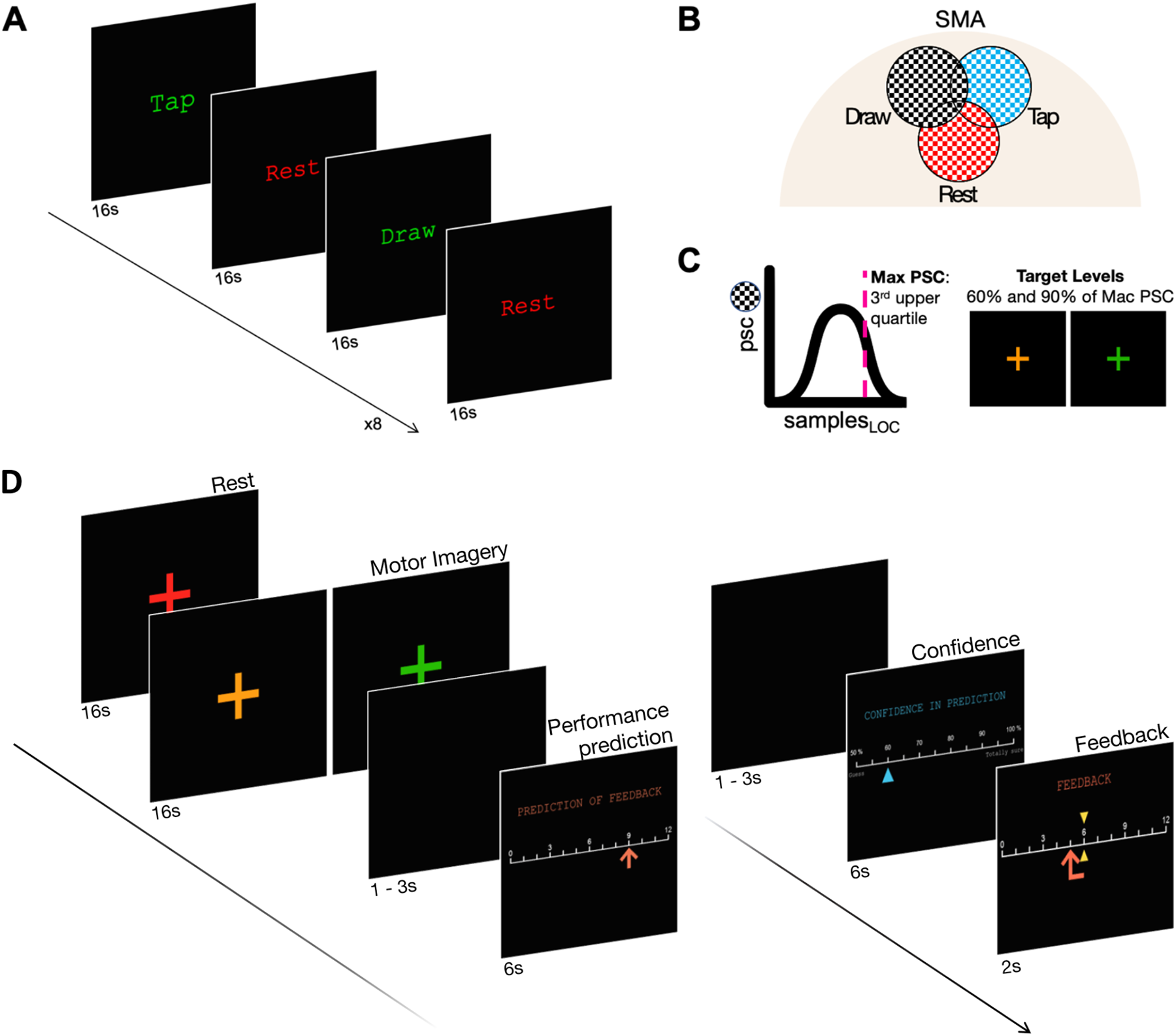
(**A**) Functional localizer task. Participants performed this task at the beginning of every session. During *Tap* blocks, participants executed real movements. During *Draw* blocks, participants imagined drawing movements. (**B**) ROI selection procedure. A target region for neurofeedback was selected by combining expert knowledge and an algorithm to select a cluster of voxels that was more active for mental drawing than for real movement and rest (see 2.5.2). (**C**) Maximum percent signal change (MaxPSC) and Target Level selection: Percent signal change (PSC) values for the imagined drawing blocks from the selected target region were used to define MaxPSC by taking the third upper quartile value. Two target levels for the neurofeedback task were defined as the 60% and 90% of the MaxPSC and were indicated by an orange or green cross, respectively. (**D**) The motor imagery neurofeedback task including self-ratings of the (1) performed regulation level (‘performance prediction’) and the (2) confidence of this performance prediction (‘confidence in prediction’), followed by the feedback of the achieved regulation level (orange arrow) and its difference to the target level (yellow marker).

##### Instructions for ratings

For self-ratings of the performed regulation level (performance prediction), participants were asked to rate the average level of activation that they thought they had achieved during the imagery period (or, in other words, to predict the neurofeedback value they would obtain for the current trial). For confidence ratings, participants were asked to rate their confidence in their predicted performance. They were reminded not to confuse this with confidence in having reached the target level or having performed better or worse. Here, we insisted that the question was about the confidence in the performance prediction (for example, they could have high confidence in a prediction that their achieved activation level would be lower than the target, or low confidence in their prediction that the neurofeedback value would be on target).

##### Control trials

In each neurofeedback run, one of ten trials was a catch trial in which the responses for the performance prediction and confidence were instructed. In these control trials, two red bars surrounded one of the values on the scale and participants were asked to move the cursor to the indicated location. These catch trials were included as an attention check, but also to reduce the confounding effect of the motor movement in the judgement process (Fleming, Huijgen, & Dolan, 2012) during offline analysis.

### 2.4 Data Acquisition

MR images were recorded using a Siemens Magnetom 7T MR scanner with a 32-channel head coil (Nova Medical Inc., Wilmington, MA, USA). High-resolution sagittal anatomical images were acquired with a T1-weighted magnetization prepared rapid acquisition gradient echo sequence (MP2RAGE; 256 sagittal slices, voxel size = 0.9 × 0.9 × 0.9 mm^3^). Functional images were obtained using a gradient echo (T2* weighted) echo-planar imaging (EPI) sequence, with the following parameters: echo time (TE) = 21 ms, repetition time (TR) = 1000 ms, multi-band factor = 3, flip angle = 60°, matrix = 224 × 224, number of slices = 60, voxel size = 2 × 2 × 2 mm^3^. The field of view provided almost whole-brain coverage.

Pulse data was recorded using a peripheral pulse unit (PPU) by Siemens, attached to the participant’s left index finger. Respiration was monitored by placing a respiratory cushion between the upper abdomen of the participant and a belt. This cushion was connected via a pressure hose to a physiologic ECG and respiratory unit (PERU) by Siemens.

Behavioral responses were recorded with a fiber optic four-button response box (Current Designs, https://www.curdes.com) attached to the participants’ right hand with the index and middle finger placed on buttons 1 and 2, respectively.

### 2.5 Online analysis

#### 2.5.1 Preprocessing

MR images were reconstructed in real time and exported to a dedicated computer, via a direct transmission control protocol / internet protocol (TCP/IP) connection, where they were preprocessed using Turbo-BrainVoyager (version 3.2, Brain Innovation B.V., Maastricht, the Netherlands). Each functional 3D volume was motion corrected to the first volume of the functional localizer run of the same session and spatially smoothed using a Gaussian kernel of 4mm. Linear trends were addressed using an additional predictor in the general linear model (GLM). The real-time processing computer and the stimulation application communicated over the network using a direct TCP/IP connection.

#### 2.5.2 Target region selection

The neurofeedback target region was defined for each participant in each session separately, combining expert knowledge and an algorithm for automated selection of brain areas (Lührs et al. 2017) based on a GLM analysis of the functional localizer data. Physiologically, the overlap between motor imagery and motor execution has been studied extensively (Hétu et al., 2013), with notable differences between M1 and the supplementary motor area (SMA). Imagined movement is better predicted by the SMA (Park et al., 2015). We therefore first used the anatomical scan to preselect an area that corresponded to the location of the SMA and then further restricted this area by using the functional localizer task (Fig 1A); the resulting area showed higher activation for mental drawing compared to finger tapping (Fig 1B for a schematic representation) and included a cluster of voxels exceeding a threshold of *t* > 5.0 for imagery vs. rest. Within this area, the algorithm automatically selected the final target region: 30 most significant voxels, forming one contiguous cluster with a 26 neighbor-voxel criterion spanning over not more than six contiguous slices. This method made it possible to select a cluster of voxels where the activation was more specific to motor imagery as compared to motor execution while also showing the highest difference between imagery and rest blocks.

#### 2.5.3 Calculation of neurofeedback

In order to provide individualized neurofeedback, the localizer’s imagery data was used to calculate MaxPSC for mental drawing blocks, for each session of each participant. Each MaxPSC was determined by calculating the third upper quartile of average mental drawing percent signal change (PSC) using a custom script in MATLAB (R2018b, MathWorks, Natick, Massachusetts). The third upper quartile was chosen instead of the maximum value to account for potential future fatigue of the participants.

Intermittent neurofeedback was then calculated as the PSC value during each mental drawing trial with respect to its preceding baseline window, adjusted for the individual session’s MaxPSC (1). To account for the BOLD delay, only the last few activation values of each rest and mental drawing period were taken into account for the PSC calculation. The baseline value corresponded to the mean activation between seconds –4 to +2 seconds after imagery block onset (6 volumes), whereas the imagery value was the average from seconds 6 to 16 (10 volumes) (see Fig. 2). The PSC of each trial was then divided by the participant’s MaxPSC and multiplied by 10 to obtain a normalized value where 10 corresponded to the MaxPSC (2).

**Figure 2.**
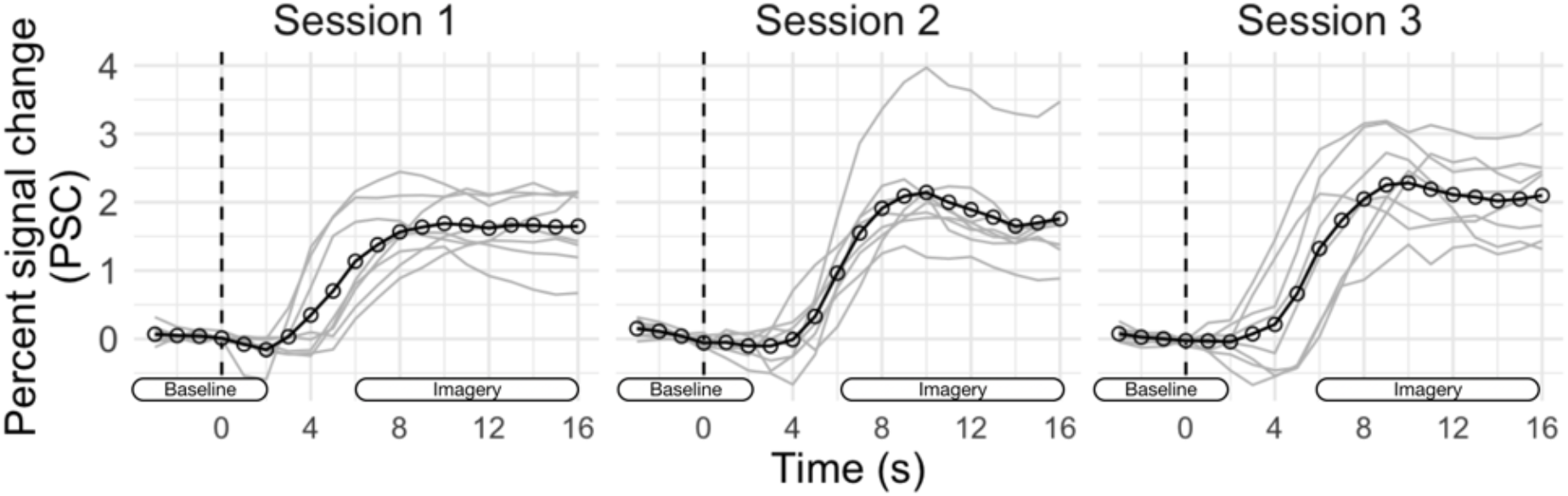
Percent signal change (PSC) for mental drawing during the functional localizer. The black lines and circles indicate the mean PSC across participants. The grey lines show each participant’s average. The text labels indicate the time ranges used for the selection of volumes for neurofeedback calculation (during neurofeedback task only). During the neurofeedback task, one functional image (volume) was acquired every second, and we ignored the first volumes of the drawing block to account for the BOLD response delay. The baseline was defined as the time period between t = –4 and t = 2 seconds with respect to mental imagery onset. The mental drawing period corresponded to the time range between t = 6 and t = 16.

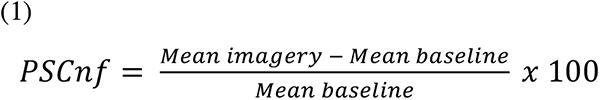

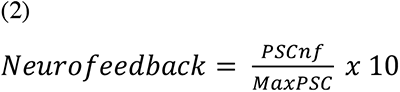

The neurofeedback and the performance prediction scale presented to the participants included values from 0 to 12, to be able to present values above the MaxPSC. Consequently, the participants were aware of a potential overshoot when regulating, which allowed them to further improve the learning process. Twelve was chosen specifically to equalize the information range between the target levels 60% and 90%, and the value presented maximally on the scale. Values below 0 and above 12 were clipped to ‘0’ and ‘12’, respectively.

### 2.6 Offline analysis

To estimate how the main outcomes (i.e., self-regulation performance, performance predictions, and confidence reports) differed according to experimental conditions, we used R (version 4.0) (R Core Team, 2020), Stan (rstan version 2.16) and the brms package (Bayesian Regression Models using Stan version 2.1.) to fit multilevel Bayesian linear models. The use of multilevel modelling allowed us to estimate the effects of interest for each participant individually (Gelman et al., 2012). The use of the Bayesian framework of *brms* over maximum likelihood-based approaches to multilevel modeling provided several benefits, such as the improved rates of convergence, the ability to make direct probability statements, and the obtention of more intuitive uncertainty estimates than those of Null-Hypothesis Significance Testing (Carpenter et al., 2017).

All three models of the main outcomes were estimated with Markov Chain Monte Carlo sampling, running 2 parallel chains for 5.000 iterations each (the first 2.000 warm-up samples for each chain were discarded). For each model we assigned random slopes and intercepts for individuals (Gelman & Hill, 2006), while priors were kept to default. We report posterior means and credible intervals (Bürkner, 2017; Carpenter et al., 2017). The posterior probability distributions from the model parameters were also used to test several hypotheses, which are listed in the subsequent sections. Since the hypothesis() function of the *brms* package does not allow to compute evidence ratios when using default priors, these hypotheses were formulated as one-directional. For each hypothesis, we therefore computed the posterior probability of the hypothesis against its alternative (for our one-directional hypotheses, this quantity corresponds to the proportion of the posterior probability above 0). The formulation of several of the one-directional effects was driven by the performance during real-time sessions and by preliminary results (e.g., subchapter 3.1.2 and corresponding Figure 4), so rather than a priori hypotheses, these should be seen as statements that guide the exploration and visualization of the results. Each hypothesis test was applied to each individual.

**Figure 3.**
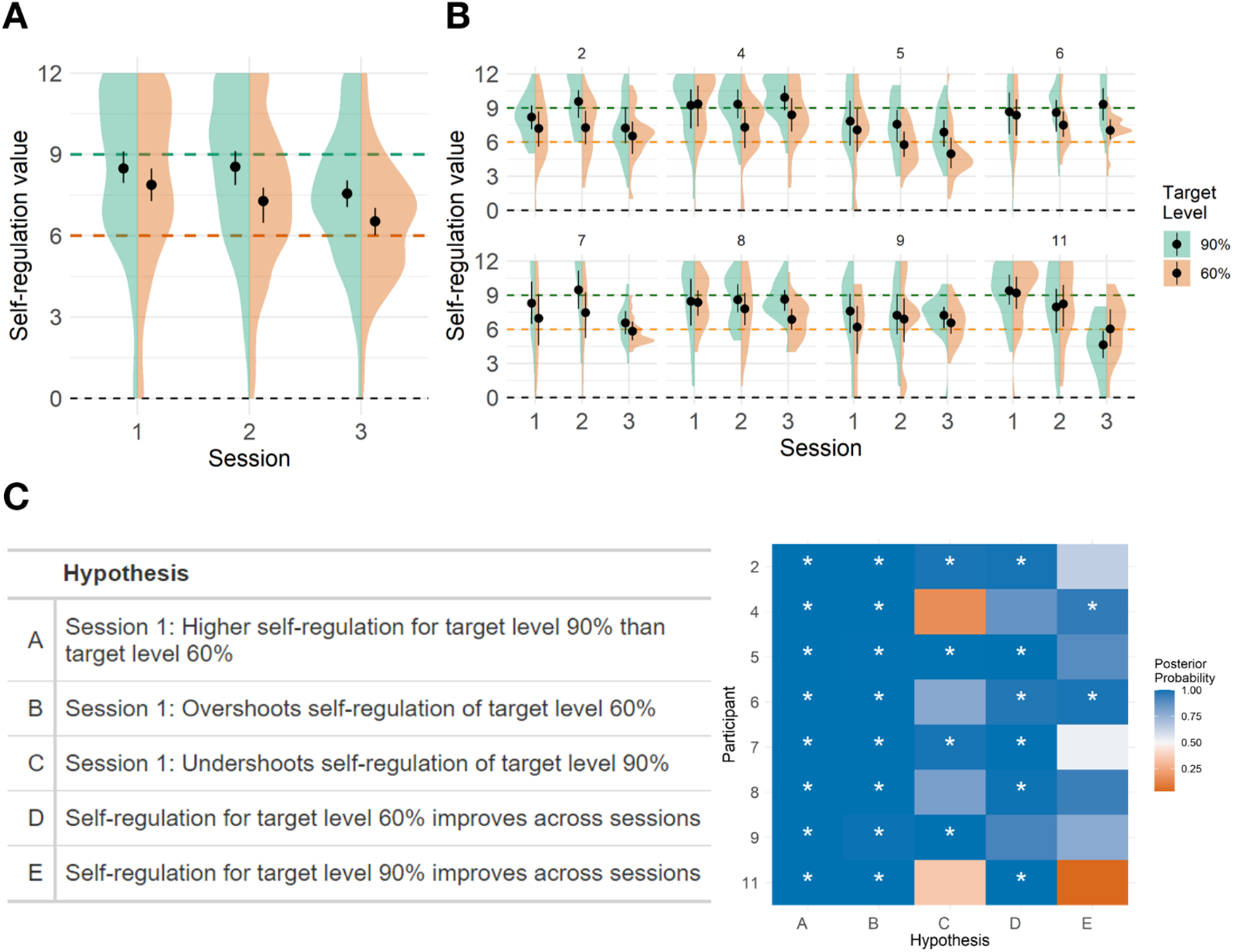
Self-regulation performance. **(A)** Group average for self-regulation performance. For each session (1, 2 and 3) and Target Level (60% or 90%, 6 or 9 on the 0-12 scale), the mean and within-subjects confidence intervals are shown. In color, the probability density distribution of the underlying data is shown, trimmed to the range of the data. **(B)** Similar to A, separated for each participant. **(C)** Table showing the posterior probability value for each hypothesis statement tested using the self-regulation model. The posterior probability corresponds to the proportion of samples from the posterior distribution of the parameters conforming to the hypothesis. A value above 0.5 (50%) indicates a higher proportion of samples in agreement with the hypothesis and is illustrated with the fill color (from red = 0% over white = 50% to blue = 100%). Asterisks indicate a posterior probability that exceeds 95%. Note that the participant identifiers equal the initial numbering (before exclusion).

**Figure 4.**
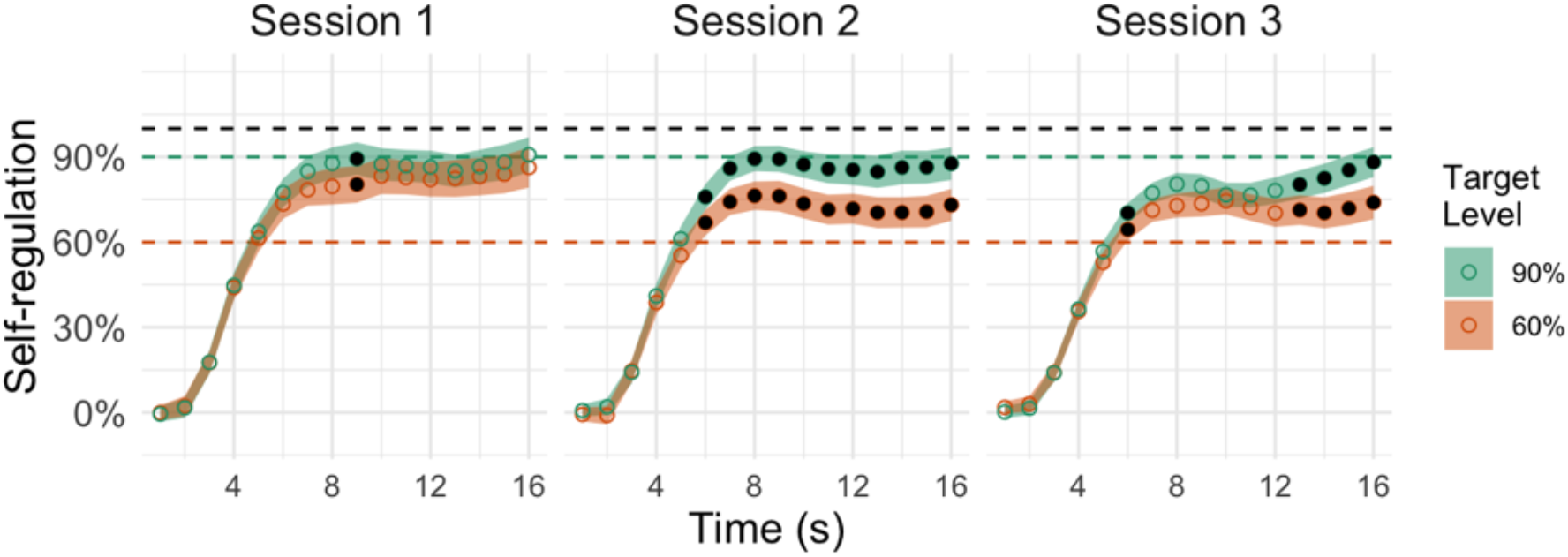
Time course of target region activation during motor imagery to two different target levels in each session. The percent signal change (PSC) is shown as a percentage of the participant’s maximum for the session (MaxPSC) as defined with the functional localizer. The horizontal black dashed line (100%) represents the MaxPSC of the session, the green dashed line (90%) represents the high Target Level (90% of the MaxPSC), and the orange dashed line (60%) represents the low Target Level (60% of MaxPSC). Each circle represents the mean for the timepoint, and colored ribbons represent a bootstrapped 95% confidence interval (CI) around the mean. Black filled dots indicate when the one-tailed t-test is significant.

For the remaining statistical analyses, null-hypothesis significance testing was used in R (version 4.0) (R Core Team, 2020) for the pointwise t-tests in the within-trial percent signal change time-course analysis (2.6.1) and MATLAB (R2018b, MathWorks, Natick, Massachusetts) for ANOVAs. If not stated otherwise, t-tests and ANOVAs were carried out two-sided and with the alpha threshold level of 0.05. For figures, R was again used with the ggplot2 package (v3.3.2, Wickham, 2016), as well as BrainVoyager (version 21.2, Brain Innovation B.V., the Netherlands) for representing brain maps.

#### 2.6.1 Percent-signal changes in target ROI

##### Whole-trial percent-signal change

To find out whether neurofeedback self-regulation performance (i.e., how far the achieved self-regulation deviated from the target level of 60% or 90%) improved across sessions, we modeled the self-regulation outcome (centered around the target level) with the *Target Level* (60% or 90%) and the Session (1, 2 or 3), as predictors. We tested five hypotheses. First, we asked whether participants achieved a higher activation for level 90% than for level 60%. We also checked whether participants undershot when trying to reach level 90% or overshot when trying to reach level 60%. In the final two hypotheses, we investigated whether the participants improved across sessions for either level. All five hypotheses are presented in Figure 3.

##### Within-trial percent-signal change time-course

Although the neurofeedback value shown to the participant (described above) was the average self-regulation over multiple seconds, we also measured and analyzed the entire time-course of self-regulation. For this analysis, we used the percent signal change (PSC) as calculated online at a frequency of 1 Hz, i.e., for each volume (for a total of 16 values per trial). We used pointwise t-tests (one-tailed for 90% > 60%) with correction for false discovery rate (FDR) at the individual level using the Benjamini-Hochberg (BH) procedure (Benjamini and Hochberg, 1995), to maintain FDR at 5% for each subject.

### 2.6.2 Performance prediction

We analyzed whether predictions of self-regulation performance became more accurate across sessions (i.e., the prediction moved closer to the actual achieved neurofeedback value), while also controlling for the effect that knowledge of previous performance might bias these predictions. Indeed, since we hypothesized that performance predictions could also be driven by feedback received in previous trials, we controlled for this possibility by calculating the running average of performance as the average neurofeedback obtained in the last five trials of the corresponding target level and modeled the performance prediction using *Target Reference* (Real Position vs. Prior of previous performance) and *Session* (1, 2 or 3) as predictors. Although a previous study used the running average of *all* previous trials in the experiment as a prior (Schurger et al., 2017), we selected a smaller time scale of 5 trials. We hypothesized that if the prior remains relatively constant for the whole experiment, our short running average will also remain relatively constant. But if there are variations at shorter timescales (within a session and within a run), this will only be captured using a shorter range for the running average. From the model results, we tested five hypotheses, also presented in Figure 5. We wanted to see whether the prediction error decreases across sessions. Then we tested whether the prediction is closer to the prior (values) or real achieved values for each session separately. Finally, we tested whether the distance between the prediction and real value decreased more than the distance between the prediction and prior.

**Figure 5.**
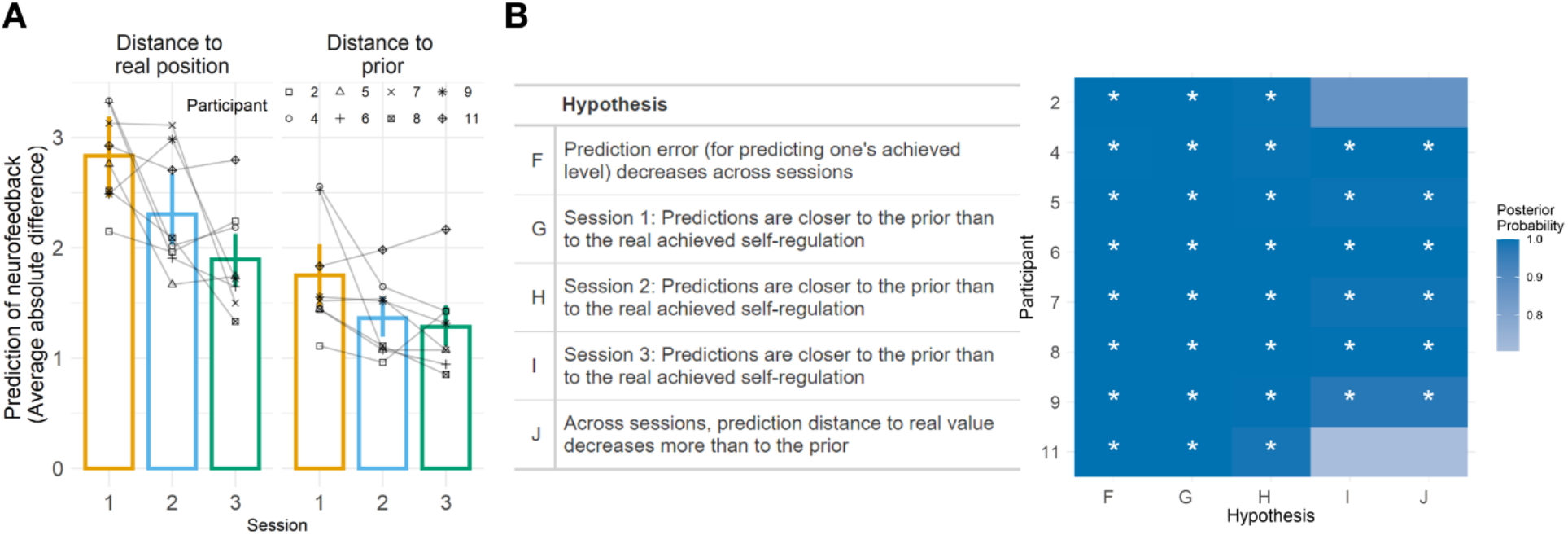
**(A)** Average performance prediction accuracy per session (in color). *Left panel*: Distance to real position (i.e., the achieved self-regulation value, as the absolute difference in each trial between the participant’s prediction and the self-regulation value). *Right panel*. Distance to the prior (i.e., the absolute difference in each trial between the participant’s prediction, and a prior consisting of the running average of self-regulation achieved in the previous 5 trials). The connected shaped dots represent each individual participant. **(B)** Individual results of hypothesis testing. Table showing the posterior probability value for each hypothesis statement tested using the performance prediction model. The posterior probability corresponds to the proportion of samples from the posterior distribution of the parameters conforming to the hypothesis. A value above 0.5 (50%) indicates a higher proportion of samples in agreement with the hypothesis and is illustrated with the fill color (from red = 0% over white = 50% to blue = 100%). Asterisks indicate a posterior probability that exceeds 95%. Note that the participant identifiers equal the initial numbering (before exclusion).

Additionally, to investigate whether better performance tends to be associated with better predictions of the actual performance, we calculated the overall and session-specific correlation between the regulation accuracy (absolute difference between target and feedback) and performance prediction accuracy (absolute difference between performance prediction and feedback).

Finally, to see if better performers (not just trials) more accurately predict their own performance, we first determined each participant’s average regulation and performance prediction accuracies (absolute differences between target levels and neurofeedback and between performance prediction and neurofeedback, respectively) for each session and then calculated a Pearson’s correlation coefficient for each session.

#### 2.6.3 Confidence in prediction

To investigate whether reported confidence changed throughout sessions and whether participants developed the ability to differentiate between their better and worse performance predictions, we modeled the confidence report on the *Performance prediction accuracy* (the absolute difference between the performance prediction and the neurofeedback value), the *Session* (1, 2 or 3), and the *Self-regulation* value (0 to 12). The self-regulation was added to account for the possibility that confidence would be influenced by the self-regulation level achieved, even though participants were asked to report their confidence that their prediction was correct. From the model results, we tested the seven hypotheses presented in Figure 6. We first wanted to find out if confidence increases over sessions. Next, we aimed at exploring whether confidence reports can be predicted by the prediction accuracy or by self-regulation accuracy. This was tested for each session separately.

**Figure 6.**
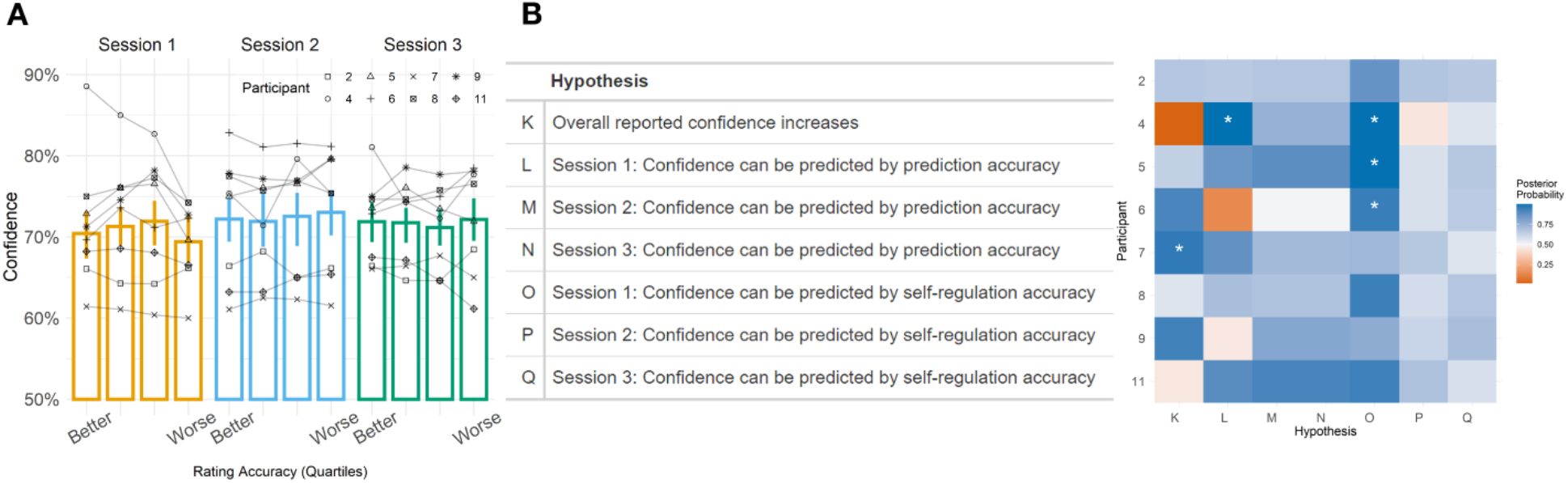
Confidence and confidence accuracy. **(A)** The graph shows trial-by-trial confidence ratings as a function of performance prediction accuracy (performance prediction accuracy – “Rating Accuracy” – has been split into quartiles for illustrative purposes only). The connected shaped dots represent each individual participant. **(B)** Individual results of hypothesis testing. Table showing the posterior probability value for each hypothesis statement tested using the confidence model. The posterior probability corresponds to the proportion of samples from the posterior distribution of the parameters conforming to the hypothesis. A value above 0.5 (50%) indicates a higher proportion of samples in agreement with the hypothesis and is illustrated with the fill color (from red = 0% over white = 50% to blue = 100%). Asterisks indicate a posterior probability that exceeds 95%. Note that the participant identifiers equal the initial numbering (before exclusion).

#### 2.6.4 Reported regulation strategies

After each run, all but one participant reported the number of mental imagery strategy changes they had made for each target level. These numbers were summed per session and level. At the end of each session, participants were also asked to represent what they were imagining drawing in a real drawing and/or description. Drawings and descriptions provided after each session were analyzed qualitatively.

#### 2.6.5 Whole-brain (f)MRI data analysis

##### Preprocessing

(f)MRI data was analyzed using BrainVoyager (version 21.2, Brain Innovation B.V., the Netherlands) and MATLAB R2018b. Anatomical data of all three sessions were first corrected for inhomogeneities, followed by the spatial transformation into ACPC space and alignment to the anatomical scan of the first MRI session. Finally, they were normalized into Talairach space.

Functional data were first preprocessed using motion correction (aligning runs to the first volume of the first run in the session), slice scan-time correction and linear trend removal, followed by the temporal high-pass filtering with four cycles per time course. The data were then corrected for EPI distortions using acquisitions with inverted phase encoding direction and the COPE plugin in BrainVoyager (Breman et al., 2020). All functional runs were aligned to the anatomical scan of the first session and normalized into Talairach space. Finally, a Gaussian spatial smoothing kernel of 6mm was applied to each run of the functional data.

##### Analysis

A fixed-effects general linear model including 7 conditions: level 60% regulation, level 90% regulation, performance prediction, confidence, performance prediction catch, confidence catch, and neurofeedback. Z-transformed motion predictors were also fitted to each participant’s data as confounds.

Maps were created for three individual contrasts. First, the two target levels (60% and 90%) were contrasted against baseline to investigate the effects of the regulation task. Second, performance prediction interval was compared to the catch trial of the performance prediction task (Performance prediction > Performance prediction catch) to find regions involved in the self-evaluation of regulation performance. Finally, the Confidence interval was compared to the catch trial of the Confidence condition (Confidence > Confidence Catch) to find regions that correspond to the self-evaluation of participants’ certainty in their prediction.

The resulting t-maps were FDR-corrected (*q* = 0.05) for multiple comparisons and three probabilistic maps were created (one for each contrast) using the individual maps of all 8 participants. Only the regions activated in all participants are reported.

## 3 Results

### 3.1 Percent signal change in target ROI

#### 3.1.1 Trial-by-trial self-regulation

In the following section, group results are described by the group posterior means (on the 0 – 12 self-regulation scale) with their 95% confidence interval. In the first session, on average, participants achieved a 0.61 [0.16 – 1.06] higher self-regulation for target level (TL) 90% than TL 60%. At the individual level, this was the case for all participants (*Hypothesis A*). In the first session, for TL 60%, participants reached an average of 7.86 [7.2 – 8.55], which was above the target. At the individual level, all participants overshot the target value (*Hypothesis B*). In the first session, for TL 90%, participants reached an average of 8.47 [7.91 – 9.01], which was below the target. At the individual level, four participants undershot compared to the target value (*Hypothesis C*). With regard to learning (difference between session 1 and session 3), results show that, on average, self-regulation for Target Level 60% improved, as the average self-regulation decreased for 1.32 [0.58 – 2.06], from 7.86 to 6.54, and therefore moved closer to the target level of 60%. At the individual level, the improvement was visible for six out of eight participants *(Hypothesis D)*. Learning effects for TL 90%, however, were less clear, as the average regulation increased by 0.42 [-0.57 – 1.42] (the range includes 0) and therefore moved closer to the target level of 90%, but individually, the improvement was only noticeable for two participants *(Hypothesis E)*. See Supplementary materials for the model summary and the list of hypothesis formulae.

#### 3.1.2 Within-trial time courses

##### Group level

We investigated the percent signal change for individual timepoints within a trial for each session at the group level. In the first session, one time point showed a significant difference between the two levels in the hypothesized direction (t = 9). In the second session, 11 timepoints were significant (t = 6 → 16). In the third session, 5 time points were significant (t = 6 and t = 13 → 16) (see **Fig 4**).

##### Individual level

A similar analysis was performed at the individual level. As shown in Figure S3 of the Supplementary materials, 75% (6/8) of the participants showed time points with significant differences between target level 60% and 90% in at least one session. In session 1, only one participant showed significant differences (P2). In session 2, three participants showed differences (P2, P4, P5). In session 3, five participants showed significant time point differences (P2, P5, P6, P8, P9) (see Figure S3 in Supplementary materials).

### 3.2 Predictions of performance

In the following section, group results are described by the group posterior means (on the 0 – 12 neurofeedback scale) with their 95% confidence interval. The results from the model show that prediction error (i.e., the absolute difference between the prediction and achieved level) decreases across sessions. On average, participants’ prediction error was 2.81 [2.45 – 3.16] in session 1, and 1.87 [1.39 – 2.35] in session 3, meaning that the prediction became 0.94 [0.46 – 1.41] points closer to the actual achieved value. At the individual level, all participants showed a decrease in prediction error across sessions (*Hypothesis F*). When looking at the potential influence of knowledge of previous performance, results show that, for all sessions, participants’ predictions are closer to the running average of previous performance than to the real achieved self-regulation in the trial. At the individual level, this was the case for all participants except for two in the last session (*Hypotheses G, H and I*). We also investigated whether previous performance could explain the improvements in performance prediction accuracy: while prediction error with respect to the prior also decreased, the decrease with respect to the real position was higher, indicating that participants’ performance predictions became closer to the real position and that changes in prior do not account for the difference. The distance to real position decreased by 0.16 [-0.31 – 0.62] in session 2, and by 0.53 [0.88 – 0.17] in session 3 more than did the distance to the prior (*Hypothesis J*). See Supplementary materials for the model summary and the list of hypothesis formulae.

#### Performance prediction accuracy and regulation accuracy

Next, the relationship between prediction accuracy and regulation accuracy was investigated. Here, a strong overall correlation of 0.81 (*P* < 0.001) was observed, with the second session showing the highest correlation of 0.86 (*P* < 0.001) between the two measures. The correlations were 0.75 (*P* < 0.001) for session 1 and 0.83 (*P* < 0.001) for session 3. Additionally, and similar to the previous step, the relationship between the prediction and regulation accuracy was investigated at the single-subject level. More accurate performers (i.e., regulating closer to the target level) indeed seem to also be better at predicting their performance (across all sessions: *r* = 0.90, *P* < 0.005 and this effect becomes more pronounced with training, specifically with sessions 2 and 3 (session 1: *r* = 0.58, *P* = 0.13; session 2: *r* = 0.94, *P* < 0.001; session 3: *r* = 0.95, *P* < 0.001).

### 3.3 Confidence in predictions

In the following section, group results are described by the group posterior means (on the 50% – 100% confidence scale) with their 95% confidence interval. The results show that on average, reported confidence for performance predictions did not increase across sessions (with an average confidence of 69.95% [63.1% – 75.6%] in session 1, and confidence of 70.5% [64.1% – 76.45%] in session 3). At the individual level, indeed, an increase in confidence was only noticeable in one participant *(Hypothesis F)*. Additionally, we found that confidence did not depend on the accuracy of the predictions, in either session. That is, participants did not report different confidence levels for their best and worst predictions of performance (S1: 0.06 [-0.17 – 0.05], S2: 0.06 [-0.08 –0.19], S3: 0.03 [-0.12 – 0.19]). Looking at individual differences, only one participant in session 1 showed an effect of performance prediction accuracy on confidence, but the effect was not present in the following sessions *(Hypotheses L, M, N)*. Lastly, while confidence was affected by the self-regulation level achieved in three participants in session 1, the effect for those three disappeared in the following sessions as well, and other participants did not show any effect of self-regulation performance in either session (S1: 0.06 [0.00 –0.13], -0.01 [-0.08 –0.06], -0.02 [-0.10 – 0.07]) (Hypotheses O, P, Q). See Supplementary materials for the model summary and the list of hypothesis formulae.

### 3.4 Regulation strategy reports and changes

Participants reported most changes to their regulation strategies occurring during the first session (mean = 2.86, SD = 2.03 for level 60% and mean = 2.43, SD = 1.18 for level 90%; total number of reported changes on group level N = 37). The number of changes for the second and third session greatly declined and stabilized, with the total of 9 changes in each session (S02: level 60% mean = 0.57, SD = 0.90; mean for level 90% = 0.71, SD = 1.16; S03: level 60% mean = 0.57, SD = 1.05; mean for level 90% = 0.71, SD = 1.16). Participants also reported imagining drawing similar objects throughout the sessions. For details, please refer to Supplementary materials.

### 3.5 Whole-brain analysis

Performing the regulation task (at both target levels – 60% and 90%) expectedly resulted in the activation of the target region (SMA) beyond the defined region-of-interest extent, but also in the bilateral activation in thalamus and basal ganglia, dorsolateral prefrontal cortex (DLPFC), insula, cerebellum and parietal regions (precuneus, inferior parietal lobe, supramarginal gyrus), and inferior frontal gyrus. The resulting clusters are presented in Figure 7 and are further described in the Supplementary materials.

**Figure 7.**
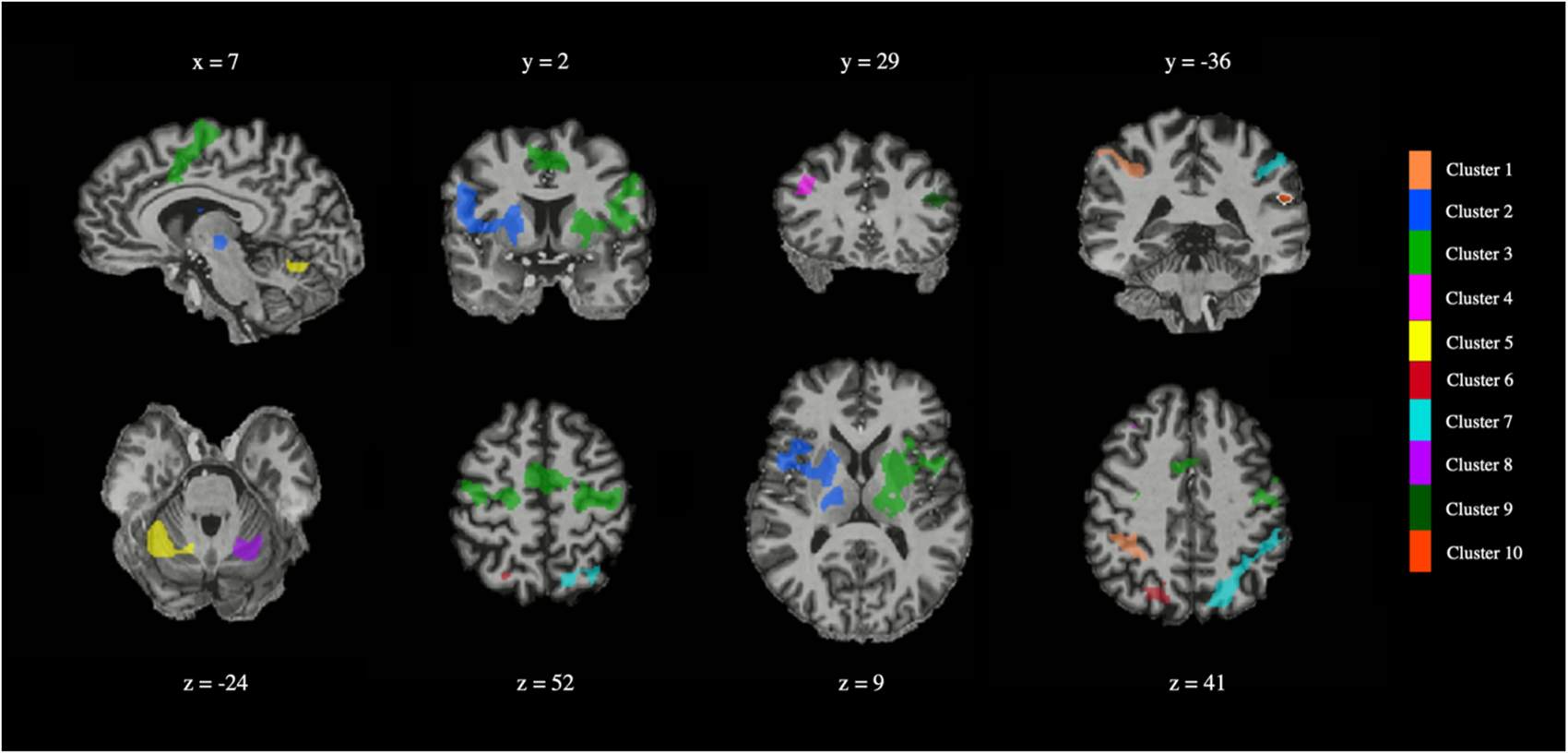
Regulation-specific activation. The two target levels (60% and 90%) were contrasted against baseline; as expected, a network related to self-regulation was revealed, together with regions involved in the task (premotor and supplementary motor areas). Each cluster is color coded and its identification number in the legend corresponds to the order of the clusters in Table S4. Note that the color-coding of clusters is merely used for visualization purposes. Brains are presented in a radiological view (i.e., right to left). The slice coordinate is written above the image in the upper row and below the image in the bottom row.

Comparing the performance prediction interval with the catch trial expectedly resulted in many activated regions, covering subcortical areas such as thalamus and basal ganglia, but also cortical areas, among others mid- and higher visual areas (cuneus, precuneus, superior parietal lobule), prefrontal regions, insular cortex, and cerebellum (see Fig. 8).

**Figure 8.**
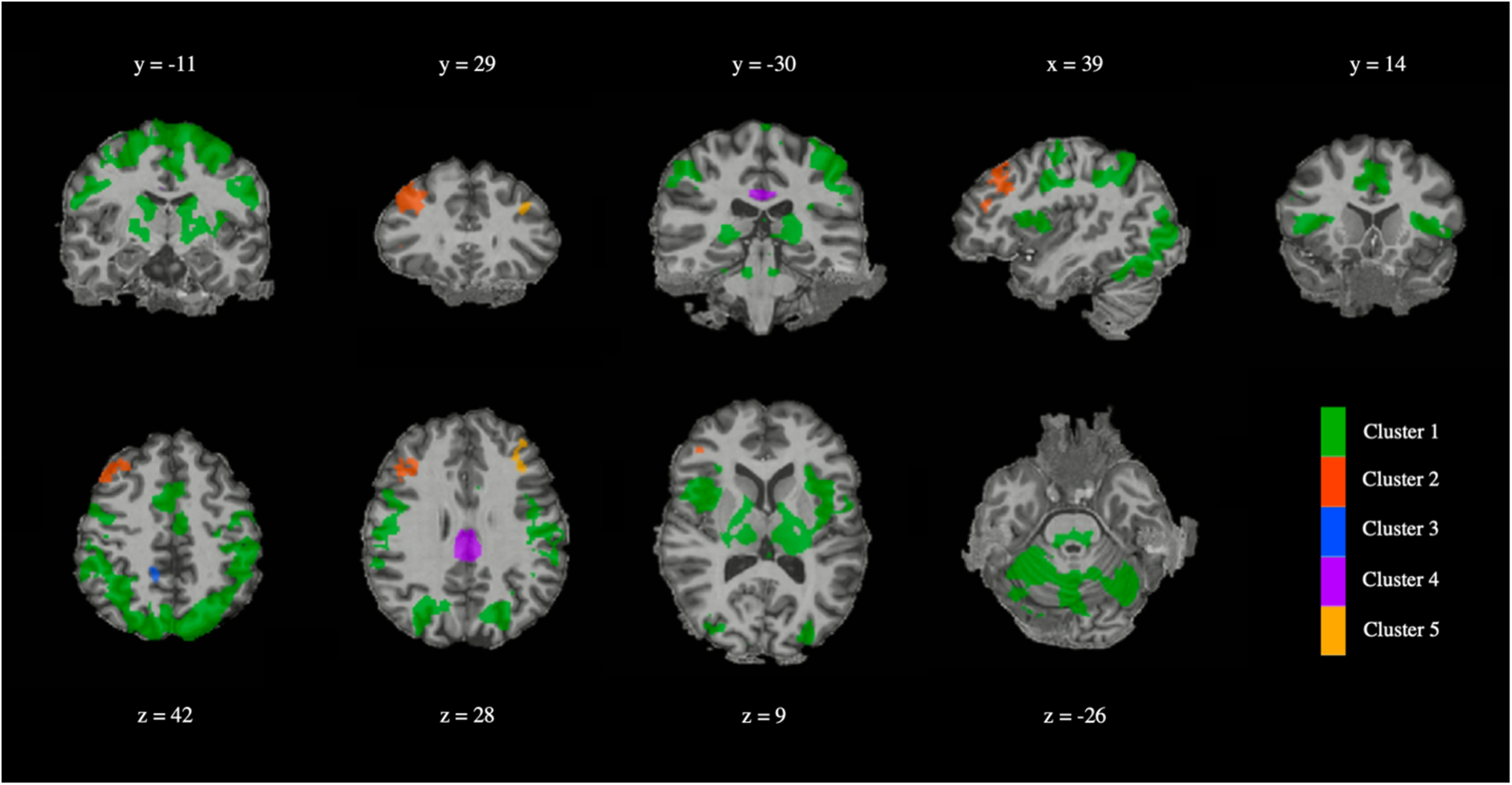
The performance prediction task was contrasted with the catch trials of the same task to control for overt movement. Both cortical and subcortical areas were activated by the task. Subcortical areas included thalamus and basal ganglia. Cortical areas involved an extensive network spreading through mid- and higher visual areas (cuneus, precuneus, superior parietal lobule), prefrontal regions (such as middle frontal gyrus) and insular cortex. Note that the color-coding of clusters is merely used for visualization purposes. Brains are presented in a radiological view (i.e., right to left). The slice coordinate is written above the image in the upper row and below the image in the bottom row.

The confidence task (confidence > confidence catch) activated most of the same areas as the performance prediction task, but to a lesser spatial extent (see Fig. 9). More detailed information is summarized in the Supplementary materials (Table S4).

**Figure 9.**
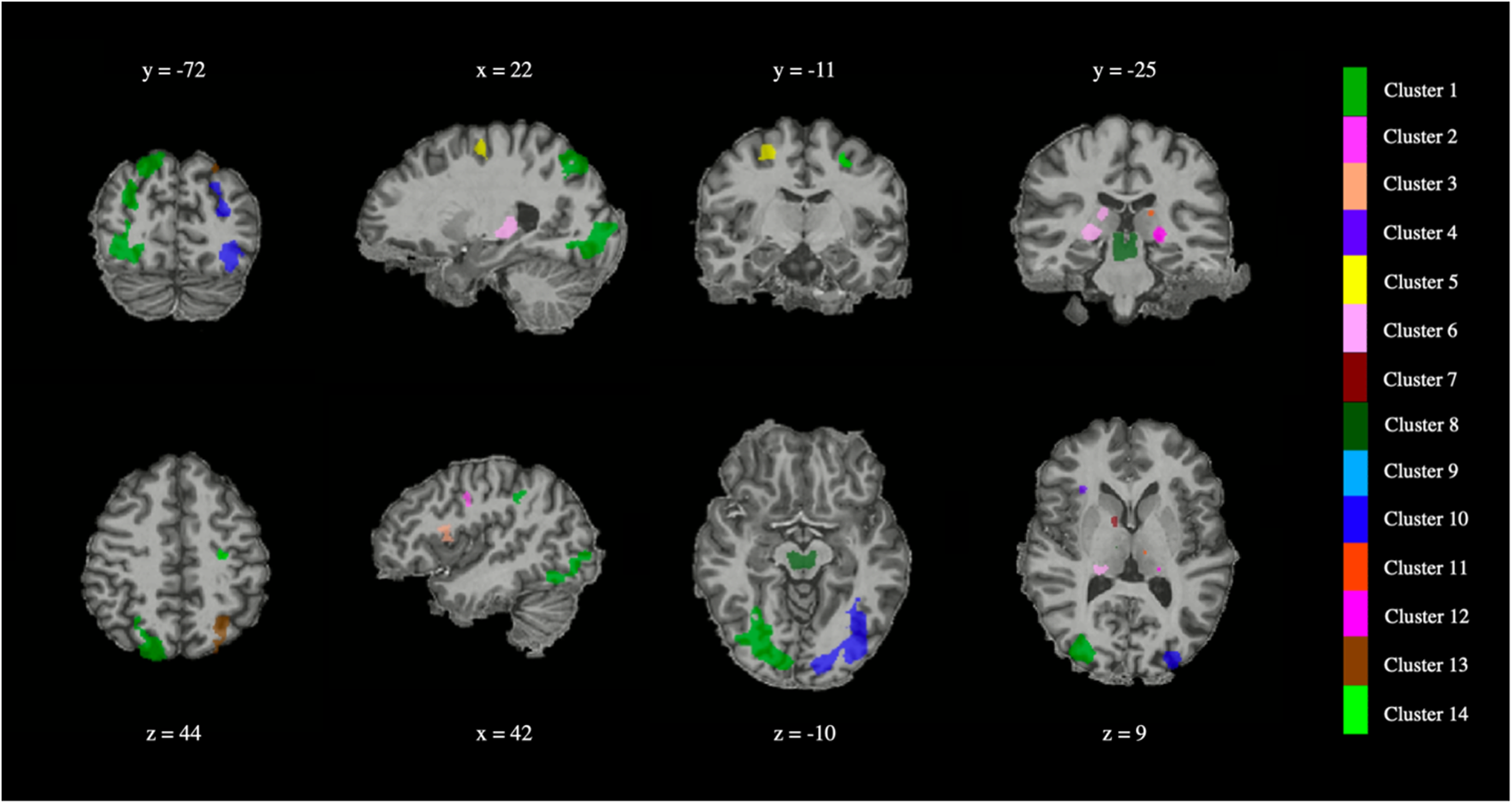
Contrasting confidence reporting trials with corresponding catch trials among others resulted in clusters covering thalamus, insular cortex, mid- and higher visual areas. Note that the color-coding of clusters is merely used for visualization purposes. Brains are presented in a radiological view (i.e., right to left). The slice coordinate is written above the image in the upper row and below the image in the bottom row.

## 4 Discussion

The capacity to monitor our ongoing mental activity is an important component of mental self-regulation, and yet its role in neurofeedback and BCI learning has remained largely unaddressed. Here, we measured people’s capacity to self-regulate the activity of a target brain region and to evaluate the level of activation they achieved and their confidence in their estimation, while receiving intermittent neurofeedback information. Intermittent feedback was crucial to obtain subjective self-reports before participants were informed about their true performance. We revealed evidence for an improvement in monitoring of mental self-regulation, confirming our hypothesis that neurofeedback guides the enhancement of self-evaluations of performance. However, the pattern of responses we observed for confidence reports invalidated our other hypothesis: although self-regulation performance and performance monitoring improved, confidence did not change and was not diagnostic of performance monitoring. We discuss the results for self-regulation, predictions of performance and confidence separately.

### Self-regulation improves with learning

Participants were asked in each trial to self-regulate their brain activity to one of the two target levels, which were adjusted for each participant based on their individual capacity. The results showed that training improved the participants’ ability to self-regulate to different target levels, which is in agreement with previous studies (Krause et al., 2017; Mehler et al., 2019; Sorger et al., 2018). Furthermore, when looking at individual data points within the entire imagery period, we were able to identify with finer detail the timepoints where significant differences between regulation to the two target levels occurred. The difference between levels was barely visible in the first session (i.e., a single time point), but it improved and reached its peak in session two with eleven time points. Fine-tuning performance then diminished in session three while still staying higher than in session one.

One interesting aspect of our results is that self-regulation performance (the ability to achieve the two separate target levels) did not improve linearly throughout the sessions, as overall performance increased in session two but decreased from session two to three. We speculate that this might be driven by changes in self-regulation strategy or motivation. Indeed, one possibility is that after the initial improvements, further progress becomes more difficult to achieve, and participants might have been more liberal in trying out other strategies, which in turn led to the performance decrease. A new strategy during the neurofeedback runs might, for example, be more difficult to finetune or could activate slightly different voxels than initially determined in the (relatively small and therefore very specific) ROI, which could in turn make the region control more difficult. Based on the participants’ reports, this however does not seem likely, as all participants reported drawing the same objects per level in the second and third session; the number of reported strategy switches also dropped after the first session and did not differentiate between the final two sessions.

Another possibility for the performance drop in the last session is that, given the attentional and cognitive load costs of sustaining self-regulation for long periods, participants’ motivation decreased when further improvements became harder to achieve. Although motivation reports were not collected, hence preventing definite conclusions, a potential solution to control for motivation in future research is to associate performance to a monetary reward.

In addition, the two target levels were not necessarily equally difficult. Given that target levels were determined based on the performance during the functional localizer (the MaxPSC, 100% on the scale), participants started with some knowledge of what 100% meant but had to learn by themselves how to obtain a level 90% or a level 60%. Level 90% is closer to 100% than level 60%, so it requires little adjustment to the strategy, which might lead to differences in difficulty. On the other hand, decreasing the signal strength by 10% to reach level 90% requires a more fine-tuned control than reaching a level 60%. Results indeed show that the average achieved signals for the two target levels in the first session were harder to differentiate and were both bellow and closer to the MaxPSC. With practice however, the differences between the two target levels became more pronounced. It has indeed been suggested that parametric training leads to steeper learning curves, as it engages participants in exploring more ways in which targeted regional activity can be self-regulated, hence accelerating the understanding of the self-regulatory process (Sorger et al., 2018).

Another explanation of the differences in difficulty to achieve the two levels might lie in the session-specific determination of the ROI region and MaxPSC value. Although the refinement of the target region proved to be advantageous (see Supplementary materials), as a higher PSC appears to facilitate the accuracy of level regulation (Krause et al., 2017), that also meant that the finetuned details of each level’s strategy changed with each session. Reaching 60% perhaps required more adjustments to the strategy than for 90%, even if kept consistent over sessions. On the other hand, redefining MaxPSC could have caused undershooting for level 90% when participants kept the same strategies over sessions. As a result of the redefinition of the target region, the chosen voxels also covered a slightly different anatomical area. The shift could potentially make it difficult to use the same strategies as in previous sessions, as the new region was possibly not (as) involved in the chosen strategy.

### Participants improve their performance prediction

After each trial, participants were asked to evaluate their performance by providing a performance prediction for that trial. We found that although participants estimated their performance more accurately when they also performed better (i.e., their regulation performance more closely matched the actual target level), they did not necessarily rely solely on their self-monitoring. Crucially, participants’ performance predictions improved throughout training, although they remained closer to their previous performance than to the real achieved values in each trial. Nevertheless, the heuristics of previous performance are not sufficient to explain all improvements in prediction accuracy.

Evaluating performance in the context of neurofeedback can be particularly difficult. In many studies of performance monitoring or metacognition, the object of evaluation is typically a form of exogenously evoked signal (e.g., as in confidence in visual perception) and is often accompanied by motor signals. People hence have access to several multi-sensory cues (sensory, motor, etc.) that can be integrated to inform their monitoring of performance (Filevich et al., 2019; Wokke et al., 2019). Because here the signal to be evaluated was self-generated, somatosensory afferents were absent or irrelevant. We speculate that this aspect inevitably led participants to use heuristics for their performance monitoring (Benwell et al., 2019; Schurger et al., 2017). As participants performed multiple trials, a heuristic for their estimates became their previous performance. We found that as participants’ predictions became more accurate with learning, the use of the heuristic diminished. Future studies could take advantage of using transfer runs or SHAM groups to investigate the effects of this heuristic on the performance prediction by not providing any feedback or incorrect feedback, respectively.

Our conclusion is in agreement with Schurger and colleagues (2017), who found that with training participants learned to better evaluate their actions performed with EEG-based motor imagery task. They are also in line with previous EEG-based neurofeedback studies looking at self-discrimination of the alpha rhythm (Frederick, 2012; Frederick et al., 2016; Kamiya, 1962). Here, using fMRI-NF, we further show that the monitoring capacity can be achieved for mental actions targeting the self-regulation of a circumscribed brain region.

### Confidence is not a reliable index of prediction accuracy

Our results provide novel data on the relation between confidence and the accuracy of performance monitoring (i.e., confidence sensitivity). Previous studies looked at the capacity to discriminate or monitor mental actions, using only evaluations of performance. Here we included an additional judgement layer, the judgement of the quality of one’s own prediction accuracy, by which we aimed to measure the participants’ ability to differentiate between simple guessing and informed judgements. We found that, although not all performance predictions were equally accurate, confidence did not differentiate between the better and the worse ones.

There are multiple ways in which confidence can relate to performance. A normative view is that confidence is a subjective probability and it is based on the probability that a choice that one made (e.g., such as a prediction in our case) was correct given the evidence (De Martino et al., 2013; Fleming et al., 2010; Meyniel et al., 2015; Pouget et al., 2016). But confidence can also be driven more directly by characteristics of the signal itself, such as its perceived uncertainty (Navajas et al., 2017) or the magnitude of sensory data (Kepecs and Mainen, 2012; Meyniel et al., 2015). For an illustration based on our task, let us imagine a given signal that a participant generated in SMA, measured in real-time as a sample of BOLD percent signal change values with mean M and variance V. An ideal observer would respond, based on our instructions for the task, as close to M as possible on the objective scale, and give a confidence report that takes into account how close (accurate) they were in their prediction (e.g., higher confidence for smaller errors). But another could more simply give a confidence estimate that depends on the uncertainty (V) of the signal, such as its inverse (1/V for the confidence scale, confidence being inversely related to the variance; lower confidence for higher variance and vice versa).

Other factors can also contribute to confidence. Here, MaxPSC was adjusted which caused an implicit adjustment of the target levels for each session; it is also unusual, if not extraordinary, for participants to obtain the desired target level more than once, making the present neurofeedback task rather difficult. Overall task difficulty, for instance, is indeed an important contributor to confidence (Festinger, 1943) and also to participants’ perceived performance ranking in the group; difficult tasks tend to make good (or experienced) performers underestimate their performance and make bad performers overestimate it (Burson et al., 2006; Kruger & Dunning, 2009). On the other hand, tasks resulting in highest performance accuracy also resulted in the lowest confidence reports, with little difference in confidence between correct and incorrect answers (Stankov and Crawford, 1997). Taking these results into consideration, we would therefore expect that the participants who predicted their performance well would rate their confidence as rather low relative to their performance; the remaining participants would misjudge their performance more, but with more confidence than their performance would suggest. The convergence of the confidence reports with training seems to be in line with this theory, especially given the difficulty of the present task and the lack of confidence improvement even when provided with feedback.

Future studies should look at how other factors affect confidence, such as difficulty, error rate in previous trials, expectancies about progress, and so on. In addition, rewarding participants for the accuracy of their confidence judgements (as in Carpenter et al., 2019) might help in achieving better confidence-performance correspondence.

### The tasks activate an extensive brain network

As expected, the regulation to the two target levels resulted in the activation of the regions related to the task itself (i.e., SMA) and self-regulation in general (such as insula, DLPFC, basal ganglia, thalamus, ACC, and inferior and superior parietal regions (Emmert et al., 2016)).

Both performance prediction and confidence self-evaluation time windows revealed clusters covering subcortical areas such as thalamus and basal ganglia, but also cortical areas, among others mid- and higher visual areas (cuneus, precuneus, superior parietal lobule), prefrontal regions (such as middle frontal gyrus), insular cortex, and cerebellum. Our primary expectation was to observe areas from a frontoparietal network previously implicated for domain-general metacognitive tasks, primarily prefrontal areas which are anatomically at the top of the cognitive hierarchy (pMPFC, vMPFC, DLPFC) and also precuneus and insula. Here, we see involvement of several regions overlapping with the ones we expected, such as left and right insula, left precuneus and inferior and middle frontal gyrus, but also many more regions not predicted. One potential caveat about the contrast analysis for self-evaluations is that we used the catch trials as a control, where the performance prediction was enforced (similar to Fleming et al., 2012b)). In our task, this included a relatively small number of control trials (n=18 catch trials vs. n=162 self-evaluation trials per participant) in order to keep the number of task-specific trials high.

Another possibility for the extensive brain activation during the two self-evaluating tasks is that there is likely still some leftover activity from the regulation trials, given the very short rest periods in between the tasks (1-3s). Longer rest periods would indeed allow the brain activity to return to baseline but would also cause the trials to become much longer, which would inevitably result in less trials per session and therefore less training. The participants would also need to keep their perceived performance in their memory for a prolonged period of time in order to correctly self-evaluate. During this time, it is possible, and indeed quite likely, that participants would think of their ratings in advance, therefore defeating the purpose of the longer rest period. Additionally, it is worth noting that self-monitoring and evaluating are a part of neurofeedback training itself and are therefore hard, if not impossible to extract.

### Potential limitations

There are several potential limiting factors to the current study. First, there is a limited number of participants (n = 8), which in principle can result in lower statistical power to detect the hypothesized effects and higher sensitivity to outliers in the sample. Nevertheless, all of our analyses were performed within-participants over several training sessions and show consistent results across participants. We also note that the use of ultra-high field (7T) fMRI is associated with improvements in the signal-to-noise ratio (up to 200%–300% when compared to 3T), thus also increasing power for statistical sensitivity (Morris et al., 2018; Torrisi et al., 2018).

Second, we were not able to measure muscular activity of the hand. Although movement was visually monitored during the initial training session outside of the scanner, it is still possible that participants relied (unconsciously) on sub-threshold muscular activity to perform self-regulation of the target region. Nevertheless, a previous study did not find that electromyographical activation was driving motor imagery (Kasahara et al., 2015). Additionally, our region-of-interest selection already partly controlled for overt movement by selecting voxels that showed higher activation for mental imagery than finger tapping.

In addition, albeit not a limitation of the study per se, the peripheral physiological parameters such as breathing and heart rate were acquired but not included in the analysis due to inconsistent data quality. Changes in these responses are associated with the magnitude of the BOLD response in the brain (Birn et al., 2008, 2006; Shmueli et al., 2007). Although none of the participants reported attentional focus to the body during self-regulation, some reported focusing on the breathing during rest periods and they could have used strategies of this kind (e.g., regulation of breathing) during the regulation periods implicitly. A previous study did indeed show slight differences in these physiological parameters with different intensities of imagery, although these were not statistically significant (Sorger et al., 2018).

Finally, to control for the knowledge of previous performance on performance predictions, we used the running average of the five previous trials. This choice was made in order to allow for the running average to reflect shorter time-scale variations in performance. However, other heuristics could potentially be used by participants, such as whole-session performance average, or even more recent performance (e.g., previous trial effects). These could also be of interest in future studies.

### Conclusions

Our results showed that participants’ performance predictions (before receiving the neurofeedback) improved throughout training, beyond what was explained by a potential heuristic based on previous performance. However, the absolute levels of confidence did not change, and the trial-by-trial confidence did not differentiate between the better and worse predictions either, indicating a dissociation between factors leading to predictions of performance and confidence.

## Supporting information

Supplementary materials - strategies and drawings

Supplementary materials - CRED-NF

Supplementary materials - extra results

## 5 CRediT author statement

**Santiago Muñoz-Moldes**: conceptualization, methodology, formal analysis, investigation, resources, data curation, writing – original draft, writing – review & editing, visualization, project administration. **Anita Tursic**: methodology, formal analysis, investigation, resources, data curation, writing – original draft, writing – review & editing, visualization, project administration. **Michael Lührs**: methodology, investigation, resources, writing – review & editing. **Judith Eck**: methodology, investigation, resources, writing – review & editing. **Amaia Benitez Andonegui**: methodology, investigation, resources, writing – review & editing. **Judith Peters**: methodology, investigation, resources, writing – review & editing. **Axel Cleeremans**: conceptualization, methodology, writing – review & editing, supervision, funding acquisition. **Rainer Goebel**: conceptualization, methodology, writing – review & editing, supervision, funding acquisition.

## 7 Acknowledgments

We would like to thank Florian Krause for his assistance with software. This work was supported by the European Commission’s Health Cooperation Work Program of the 7th Framework Program, under the Grant Agreement ‘BRAINTRAIN’, grant no. 602186, a European Research Council Advanced Grant ‘RADICAL’ to AC, grant no. 340718, and a FNRS mobility grant to SMM, grant no. 2017/V3/5/137IB/JN2110, and further financial support from the Wiener-Anspach Foundation to SMM. The funders had no role in study design, data collection and analysis, or preparation of the manuscript.

## 8 Data sharing

Experimental materials, analysis scripts, and behavioral data (including online neurofeedback values) have been made available on a permanent archive (https://osf.io/n82x7/). Neuroimaging data is available upon request to keep the privacy of research participants uncompromised.

