## Supplementary materials - strategies and drawings for "Online self-evaluation of fMRI-based neurofeedback performance"

**Strategy reports**

| **Participant** | **Level** | **Session 1** | **Session 2** | **Session 3** |
| --- | --- | --- | --- | --- |
| P02 |  | \ | \ | \ |
| P04 | 6 | 2 | 2 | 1 |
|  | 9 | 2 | 2 | 3 |
| P05 | 6 | 7 | 2 | 0 |
|  | 9 | 3 | 3 | 0 |
| P06 | 6 | 2 | 0 | 0 |
|  | 9 | 3 | 0 | 0 |
| P07 | 6 | 2 | 0 | 0 |
|  | 9 | 2 | 0 | 0 |
| P08 | 6 | 4 | 0 | 0 |
|  | 9 | 4 | 0 | 0 |
| P09 | 6 | 3 | 0 | 3 |
|  | 9 | 3 | 0 | 2 |
| P11 | 6 | 0 | 0 | 0 |
|  | 9 | 0 | 0 | 0 |
| **Total count (N)** |  |  |  |  |
| Both levels |  | 37 | 9 | 9 |
| Level 6 |  | 20 | 4 | 4 |
| Level 9 |  | 17 | 5 | 5 |
| **Mean (SD)** |  |  |  |  |
| Both levels |  | 2.64 (1.67) | 0.64 (1.04) | 0.64 (1.11) |
| Level 6 |  | 2.86 (2.03) | 0.57 (0.90) | 0.57 (1.05) |
| Level 9 |  | 2.43 (1.18) | 0.71 (1.16) | 0.71 (1.16) |

**Table S4.** Reported number of strategy changes per session for each participant and level combination.

**Participant 02**

Data not available

**Participant 04**

*Session 1*

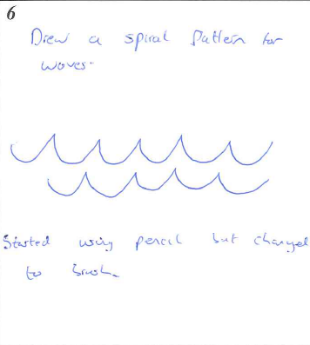

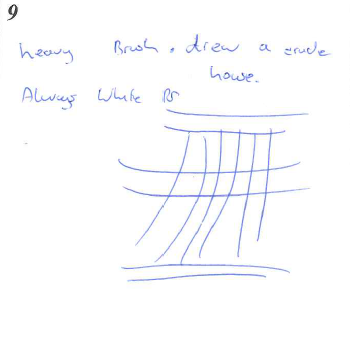

Text on figures: 6: Drew a spiral pattern for waves. Started with pencil that changed to brush. 9: Heavy brush. Drew a crude house. Always white

Comments: /

*Session 2*

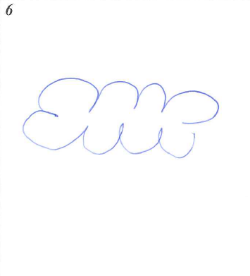

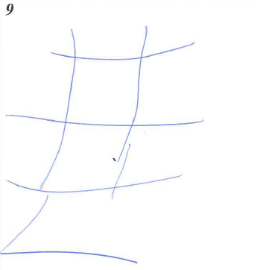

Comments: If I slowed down to reduce the score, I had to try too hard, which increased the score (i.e., neurofeedback). For one trial I inverted the drawings, drawing 6 during 9 and 9 during 6, then I switched back.

Comments: /

*Session 3*

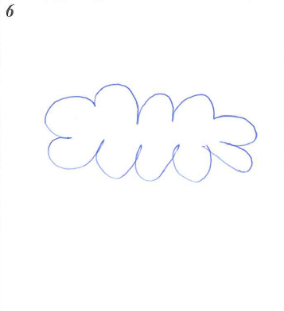

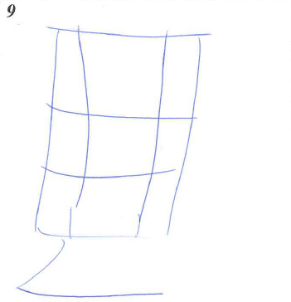

Comments: No major changes. I altered the method of drawing during level 9 at random, changing the sequence.

**Participant 05**

*Session 1*

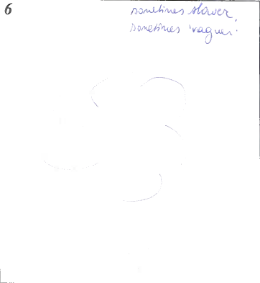

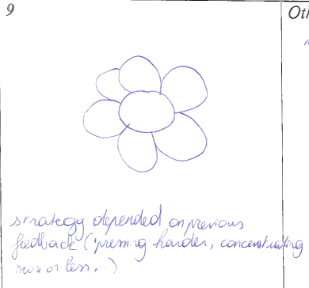

Text on figures: 6: Sometimes slower, sometimes ‘vaguer”. 9: strategy depended on previous feedback (pressing harder, concentrating more or less)

Comments: / (distracted by the scanner noise, more difficult to focus than during training outside of the scanner)

*Session 2*

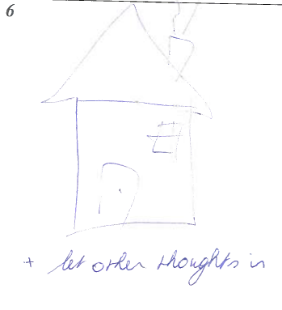

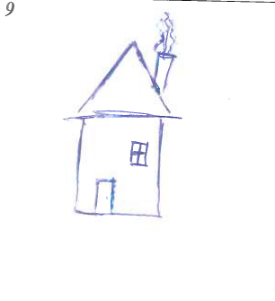

Text on figures: 6: + let other thoughts in

Comments: /

*Session 3*

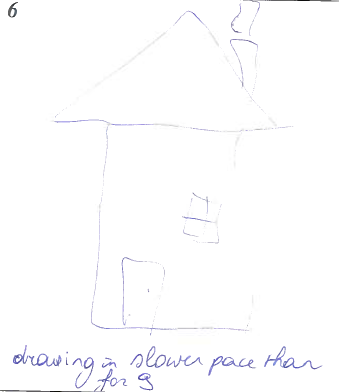

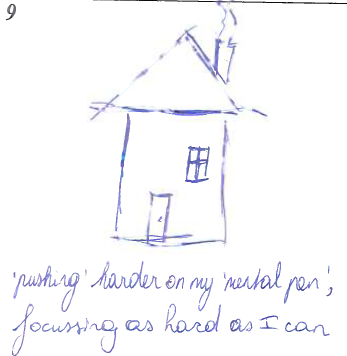

Text on figures: 6: drawing in slower pace than for 9. 9: pushing harder on my “mental pen”, focusing as hard as I can

Comments: /

**Participant 06**

*Session 1*

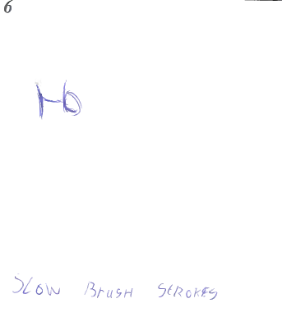

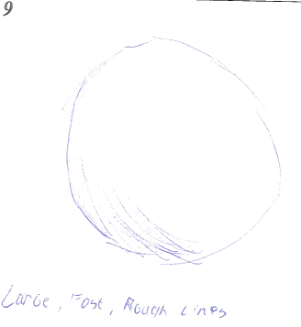

Text on figures: 6: slow brush strokes. 9: large, fast, rough lines.

Comments: /

*Session 2*

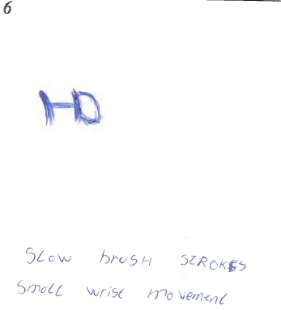

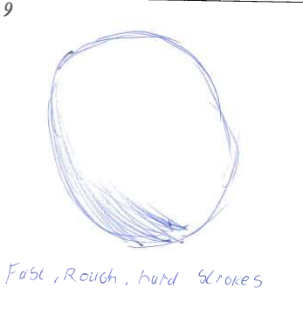

Text on figures: 6: slow brush strokes, small wrist movement. 9: fast, rough, hard strokes.

Comments: /

*Session 3*

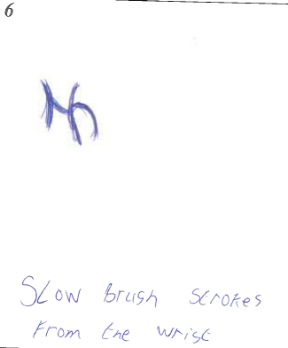

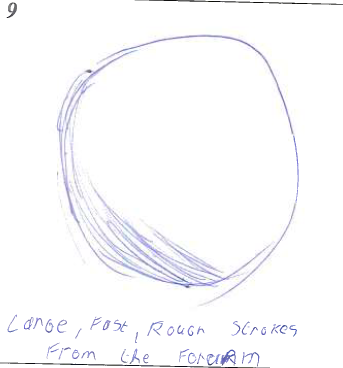

Text on figures: 6: slow brush strokes, from the wrist. 9: large, fast, rough strokes from the forearm.

Comments: /

**Participant 07**

*Session 1*

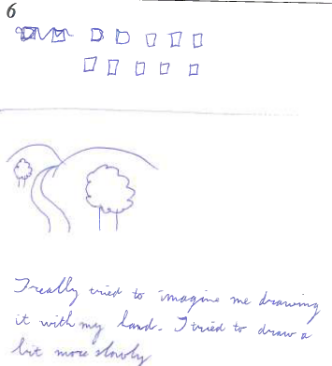

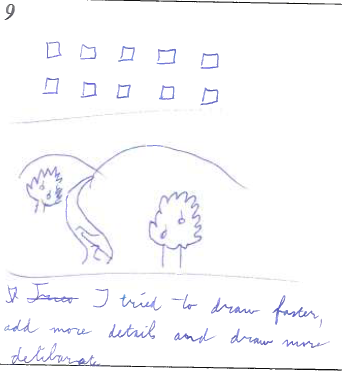

Text on figures: 6: I really tried to imagine me drawing it with my hand. I tried to draw a bit more slowly. 9: I tried to draw faster, add more details and draw more deliberate.

Comments: I tried to really imagine drawing with my hand.

*Session 2*

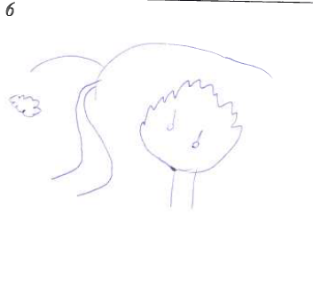

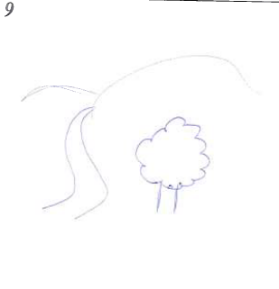

Comments: I noticed imagining my hand drawing increased neurofeedback score a lot. I omitted the imagined hand this time. I also tried to remain more bodily calm (no little twitches).

*Session 3*

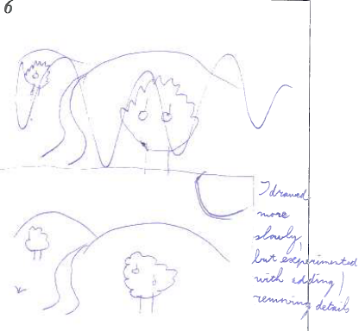

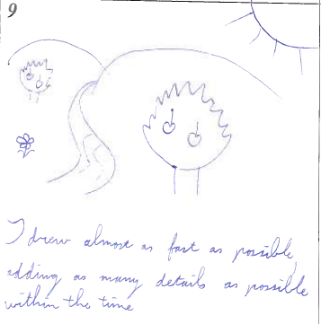

Text on figures: 6: I drew more slowly, but experimented with adding/removing details. 9: I drew almost as fast as possible, adding as many details as possible within the time.

Comments: I imagined drawing without visualizing a hand drawing.

**Participant 08**

*Session 1*

(the participant mainly provided only descriptions due to the detailed nature of the drawings, which they did not feel capable of drawing in real life)

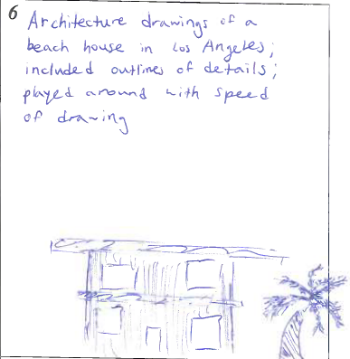

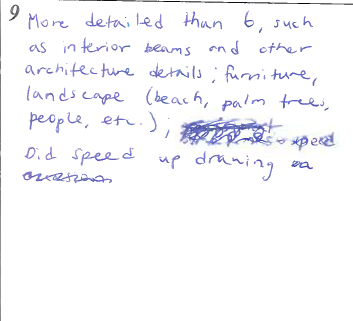

Text on figures: 6: Architecture drawings of a beach house in Los Angeles included outlines of details. Played around with speed of drawing. 9: More detailed that 6, such as interior beams and other architecture details; furniture, landscape (beach, palm trees, people, etc.). Did speed up drawing.

*Session 2*

6: Same as before, architecture drawings of a beach house in Los Angeles included outlines of details, strategies used were speed and depth in detail. Mostly played with details in this session.

9: More detail oriented, more interior, textures, lightning, etc.; also more landscape and location of environment (including smell and feel of the ocean).

*Session 3*

6: Imagined drawing a beach house in Los Angeles and walking through it while drawing. Speed and details were factors. Not as fast as 9 and details were more structural, general.

9: compared to 6, details were more prominent (textures, colors, etc.). on occasion sped up drawing, but mostly reverted to details. When neurofeedback seemed low, attempted to incorporate some senses to boost the level.

Comments: for maximum activation used senses (smell of salt water, feeling of the ocean breeze, sunlight, etc.) and details of the environment.

**Participant 09**

*Session 1*

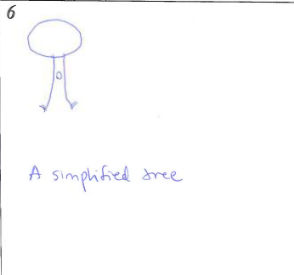

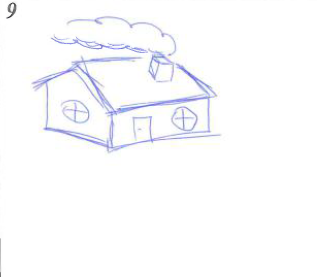

Text on figures: 6: A simplified tree.

Comments: I varied the speed of my drawing too, so that the house was faster and more details could be drawn within that timeframe. Similarly, the tree drawing was slow.

*Session 2*

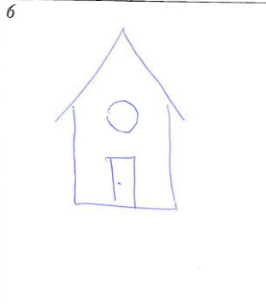

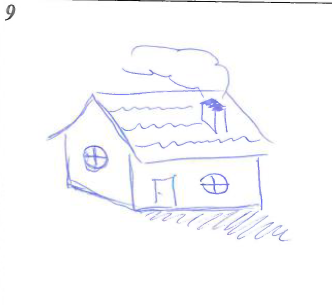

Comments: I drew faster for level 9 and tried to make more shadows for the house. For level 6 I drew slower, simpler.

*Session 3*

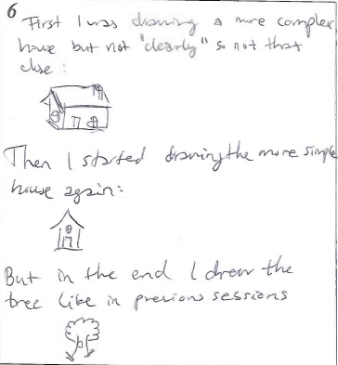

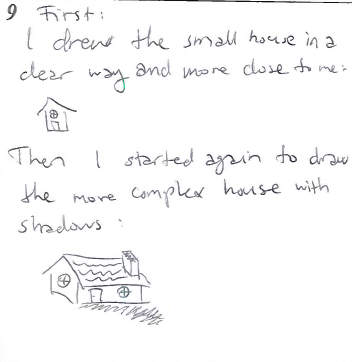

Text on figures: 6 first I was drawing a more complex house but not “clearly” so not that close. Then I stared drawing the more simple house again. But in the end I drew the tree like in previous sessions. 9: I drew the small house in a clear way and more close to me. Then I started again to draw the more complex house with shadows.

Comments: Imagining drawing more “far away” helped for level 6, and for level 9 I drew more close by. I also dried to draw smaller lines for level 6, but that didn’t really matter.

**Participant 11**

*Session 1 – 3*

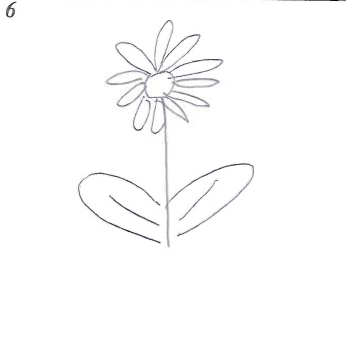

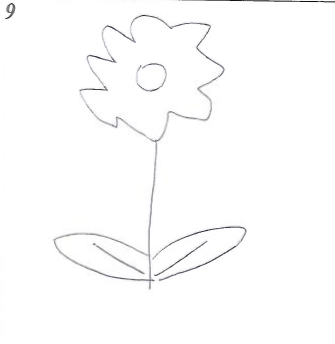

Session 3 comments: The only change was that I tried to be relaxed for rest steps.
