## Supplementary materials - CRED-NF for "Online self-evaluation of fMRI-based neurofeedback performance"

**Consensus on the Reporting and Experimental Design of clinical and cognitive-behavioural Neurofeedback studies (CRED-nf) best practices checklist 2020*** (an online tool to complete this checklist is available at [[rtfin.org/CREDnf](http://www.rtfin.org/CREDnf)](http://www.rtfin.org/CREDnf)).

| **Domain** | **Item #** | **Checklist item** | **Reported on page #** |
| --- | --- | --- | --- |
| **Pre-experiment** | | | |
|  | 1a | Pre-register experimental protocol and planned analyses | x |
|  | 1b | Justify sample size | x |
| **Control groups** | | | |
|  | 2a | Employ control group(s) or control condition(s) | x |
|  | 2b | When leveraging experimental designs where a double-blind is possible, use a double-blind | n.a. |
|  | 2c | Blind those who rate the outcomes, and when possible, the statisticians involved | n.a. |
|  | 2d | Examine to what extent participants and experimenters remain blinded | n.a. |
|  | 2e | In clinical efficacy studies, employ a standard-of-care intervention group as a benchmark for improvement | n.a. |
| **Control measures** | | | |
|  | 3a | Collect data on psychosocial factors | x |
|  | 3b | Report whether participants were provided with a strategy | 8 |
|  | 3c | Report the strategies participants used | Suppl. |
|  | 3d | Report methods used for online-data processing and artifact correction | 10 |
|  | 3e | Report condition and group effects for artifacts | x |
| **Feedback specifications** | | | |
|  | 4a | Report how the online-feature extraction was defined | 11 |
|  | 4b | Report and justify the reinforcement schedule | 9 |
|  | 4c | Report the feedback modality and content | 8 |
|  | 4d | Collect and report all brain activity variable(s) and/or contrasts used for feedback, as displayed to experimental participants | 11 |
|  | 4e | Report the hardware and software used | 10 |
| **Outcome measures** | | | |
| Brain | 5a | Report neurofeedback regulation success based on the feedback signal | 17 |
|  | 5b | Plot within-session and between-session regulation blocks of feedback variable(s), as well as pre-to-post resting baselines or contrasts | 18, except within-session |
|  | 5c | Statistically compare the experimental condition/group to the control condition(s)/group(s) (not only each group to baseline measures) | n.a. |
| Behaviour | 6a | Include measures of clinical or behavioural significance, defined a priori, and describe whether they were reached | 20 |
|  | 6b | Run correlational analyses between regulation success and behavioural outcomes | 20 |
| **Data storage** | | |  |
|  | 7a | Upload all materials, analysis scripts, code, and raw data used for analyses, as well as final values, to an open access data repository, when feasible | 33 |

*Darker shaded boxes represent *Essential* checklist items; lightly shaded boxes represent *Encouraged* checklist items. We recommend using this checklist in conjunction with the standardized CRED-nf online tool ([rtfin.org/CREDnf](http://www.rtfin.org/CREDnf)) and the CRED-nf article, which explains the motivation behind this checklist and provides details regarding many of the checklist items.
