## Supplementary materials - extra results for "Online self-evaluation of fMRI-based neurofeedback performance"

Supplementary information

Target region selection

**Target region coordinates per session and participant**

| **Participant** | **Session** | **Talairach coordinates (center of gravity)** | | | **MaxPSC** |
| --- | --- | --- | --- | --- | --- |
| P2 | S01 | -6 | -7 | 66 | 2.52 |
|  | S02 | -6 | -9 | 65 | 1.98 |
|  | S03 | -6 | -7 | 67 | 3.59 |
| P4 | S01 | -2 | 1 | 57 | 1.64 |
|  | S02 | -5 | -5 | 54 | 1.91 |
|  | S03 | -5 | -7 | 53 | 1.58 |
| P5 | S01 | 0 | -11 | 61 | 1.52 |
|  | S02 | -20 | -10 | 48 | 2.24 |
|  | S03 | -22 | -9 | 50 | 2.28 |
| P6 | S01 | -4 | -26 | 70 | 2.76 |
|  | S02 | -4 | -26 | 69 | 4.09 |
|  | S03 | -4 | -28 | 68 | 2.14 |
| P7 | S01 | -10 | 4 | 53 | 1.53 |
|  | S02 | -9 | 5 | 56 | 1.42 |
|  | S03 | -9 | 5 | 51 | 3.17 |
| P8 | S01 | -8 | 3 | 50 | 1.77 |
|  | S02 | -5 | 2 | 49 | 2.01 |
|  | S03 | -11 | 5 | 45 | 2.29 |
| P9 | S01 | -9 | 12 | 60 | 1.91 |
|  | S02 | -7 | 13 | 58 | 2.25 |
|  | S03 | -5 | 9 | 58 | 2.84 |
| P11 | S01 | -5 | -10 | 64 | 2.28 |
|  | S02 | -6 | -10 | 69 | 2.18 |
|  | S03 | -7 | -8 | 69 | 3.71 |

**Table S1.** Coordinates of target region selection per session and participant.

**The implication of session-specific ROI selection**

*The session-specific definition of the region-of-interest.* Although not the main interest of the study, the effect of the ROI redefinition for each participant in each session was also investigated.

In order to explore subject- and session-dependent region of interest (ROI) selection for neurofeedback and its potential effect on up-regulation performance in a motor imagery task, ROI time courses of each localizer run were extracted using MATLAB (R2018b, MathWorks, Natick, Massachusetts). To confirm that the potential differences cannot be attributed to merely a better selection of the region, the MaxPSC was calculated for each session-specific ROI in each session. The values of each ROI-session combination were compared in 3x3 repeated measures ANOVA (3 ROIs and 3 sessions) (SPSS24, IBM Corp.) to inspect the effect of individual ROI selection on participants’ performance. The performance was defined by the up-regulation of the ROI, therefore by the MaxPSC. Additionally, the Euclidean distances between the centers-of-gravity of the three session-specific ROIs were averaged across participants to determine spatial variability across sessions.

To see if redefinition of the ROI for each participant in each session contributes to their regulation performance, the ROI-session combinations were compared based on their MaxPSC values (see Fig. S1). The average PSC seemed to increase over sessions, although this was not significant. Nevertheless, comparing the activation between sessions 1 (S01) and 3 (S03) within the selected region of session 3 (ROI S03) resulted in a close to significance activation increase (p = .054, Sidak corrected). Moreover, the MaxPSC was on average the highest for the session-specific ROI. The difference between the MaxPSC when comparing different ROIs became significant in session 3, where the ROI selected in the corresponding session (S03) showed a significanlty higher activation than the regions selexted in first session (S01, p < .01, Sidak corrected) or second session (S02, p < .05, Sidak corrected). All in all, the redefinition of the target region seems to be advantageous.

Taking into account the voxel size (2x2x2 mm) and comparing mean Euclidean distances between ROIs (7.04 mm between ROI S01 and ROI S03 and 3.83 mm between ROI S02 and ROI S03), regions still partially overlap (also see Fig. S2). Considering the overlap and the automatic selection of most significant voxels, there seems to be a spatial shift in activation, which seems beneficial for self-regulation learning (based on increase of MaxPSC).

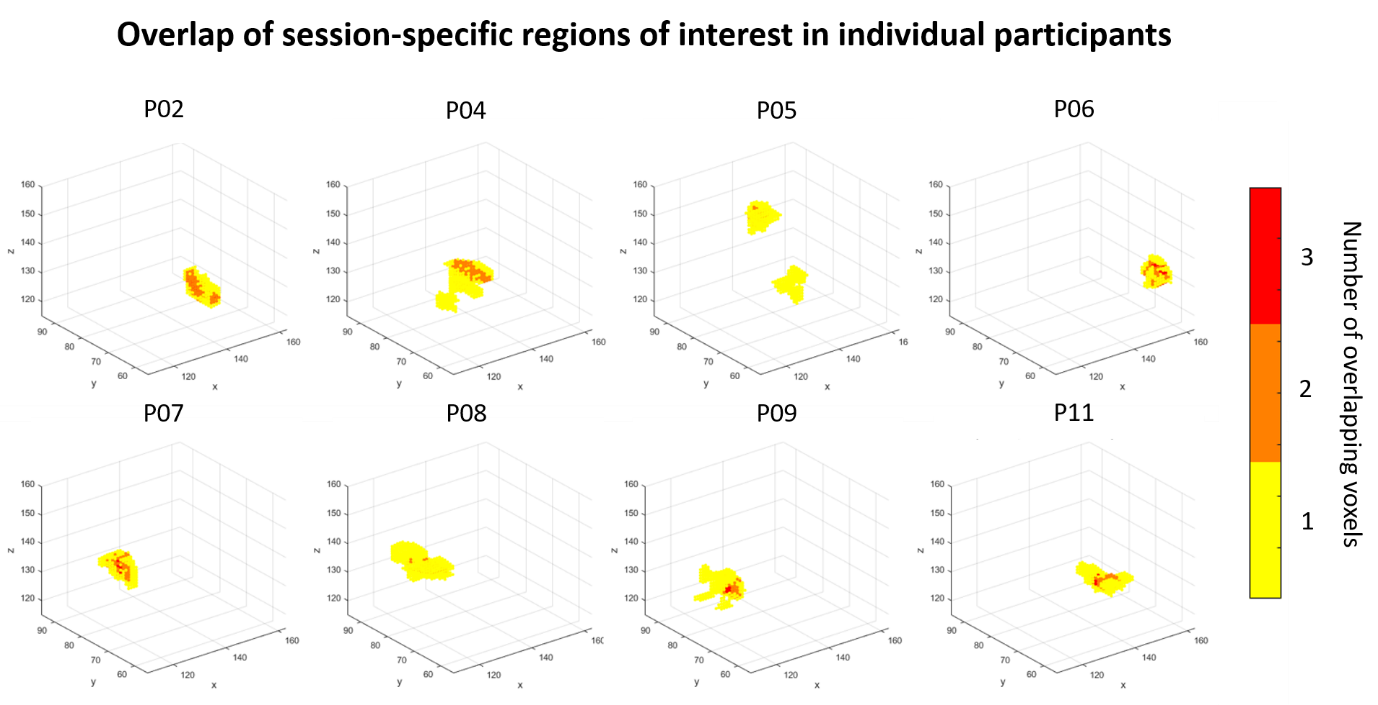

**Figure S1**. Overlap of the three regions of interest, plotted in 3D space for each participant. On the axes are coordinates in native space. Yellow color represents unique voxels of each session, and in orange and red the overlapping voxels of two or all three regions, respectively, are plotted.

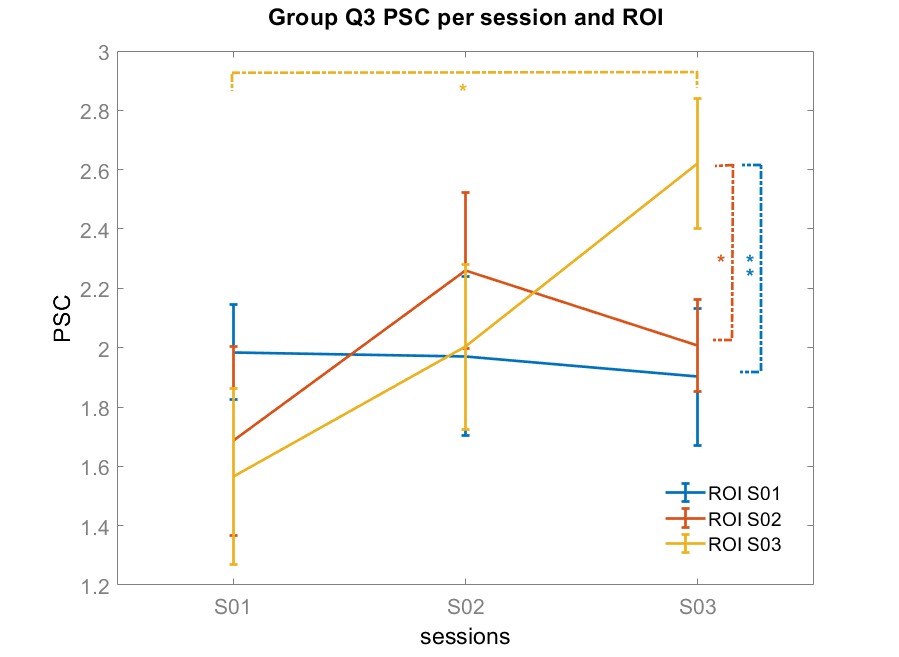

**Figure S2**. MaxPSC values for the combination of each session (S01 – S03) and region-of-interest. A significant difference in MaxPSC was observed in session 3 (S03), where the MaxPSC of the third session’s ROI (ROI S03, in yellow) was significantly higher than the MaxPSC of the other two sessions (ROI S01, in blue, and ROI S02, in orange). Comparing the first and last session’s MaxPSC in corresponding region-of-interest (ROI S03, yellow) also revealed a significant difference. Note: * p<.05, ** p<.01.

Time course of target region activation per participant

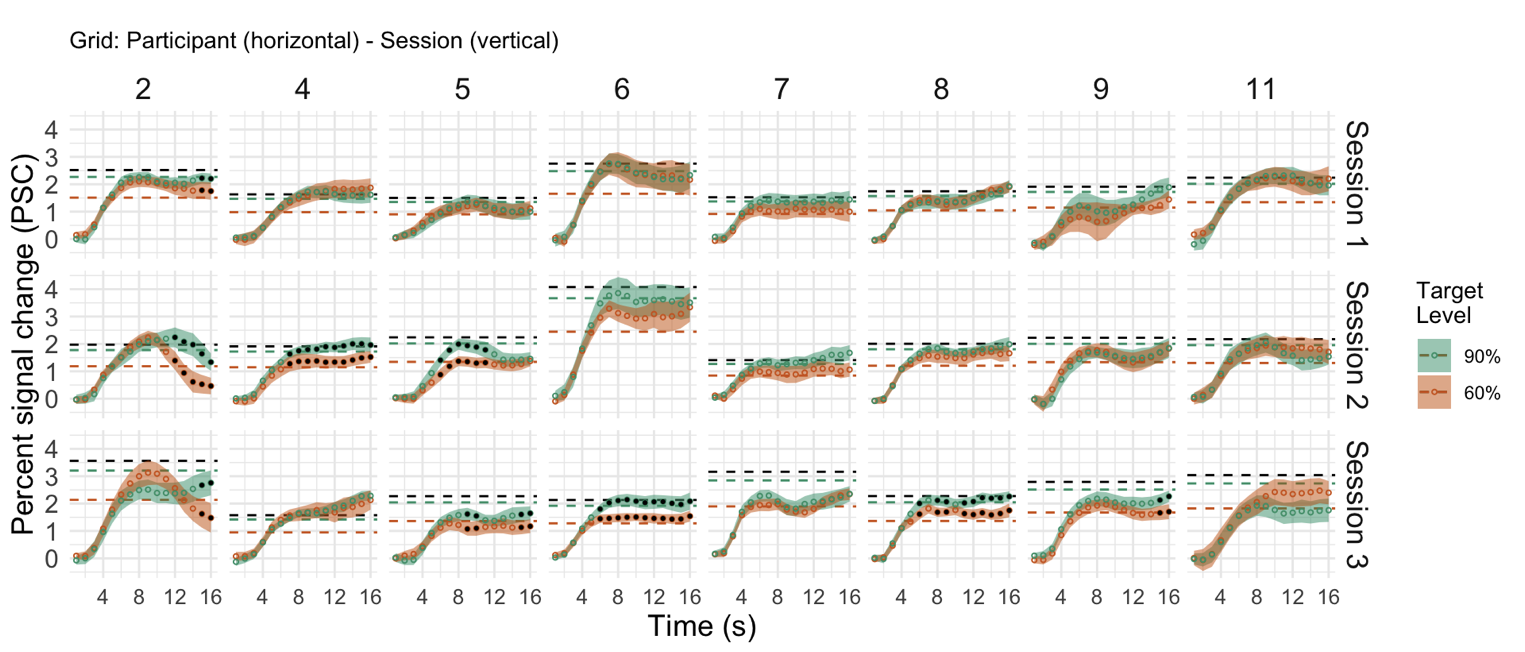

**Figure S3**. Time course of target region activation (the percent signal change; PSC) during motor imagery to two different target levels in each session for each participant. The horizontal black dashed line (100%) represents the MaxPSC of the session as defined in the functional localize. The green dashed line (90%) represents the high Target Level (90% of the MaxPSC) and the orange dashed line (60%) represents the low Target Level (60% of MaxPSC). Each circle represents the mean for the timepoint, and colored ribbons represent a bootstrapped 95% confidence interval (CI) around the mean. Black filled dots indicate when the one-tailed t-test was significant.

Hypothesis testing: models and summaries

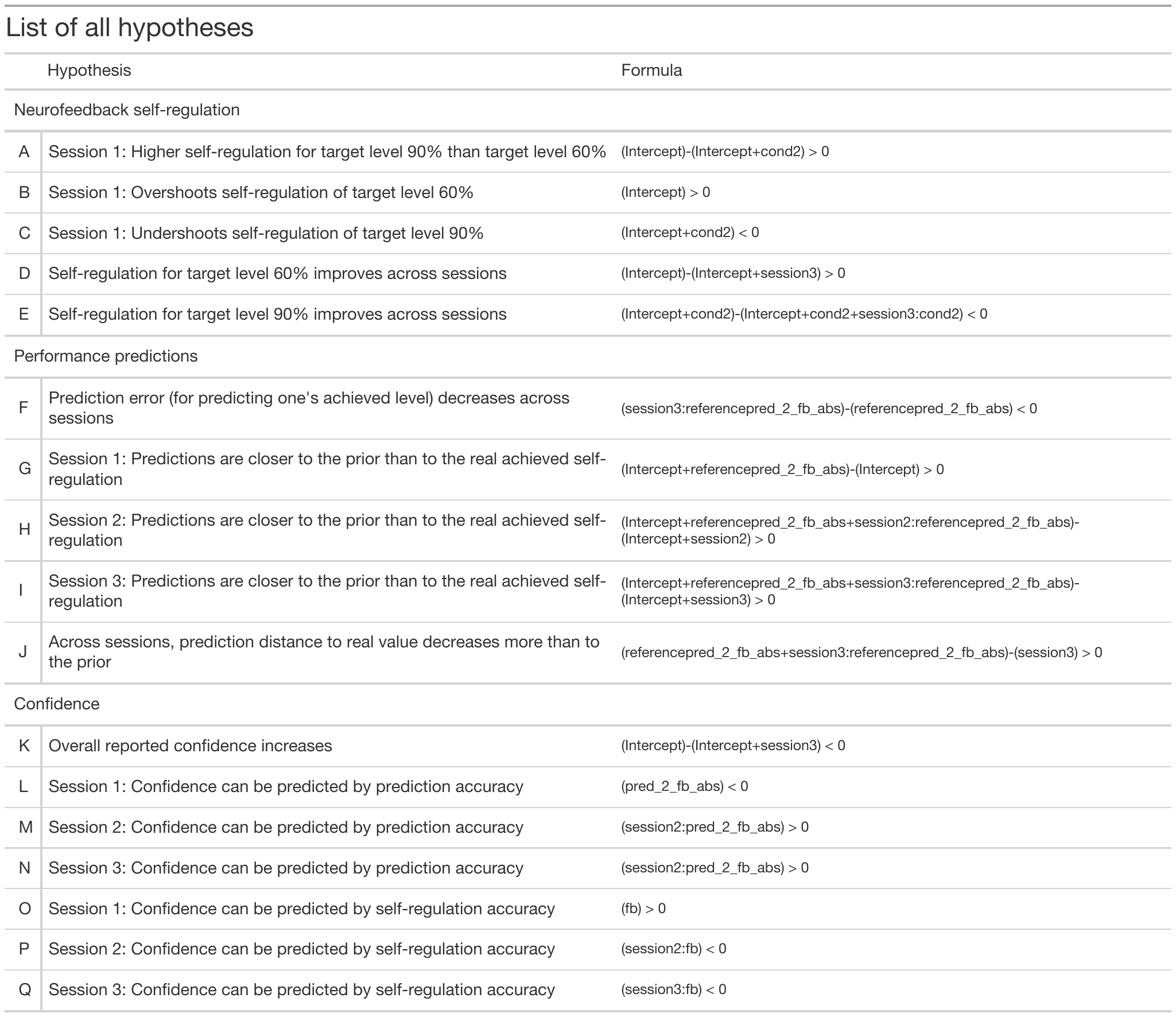

**Table S2**. List of all hypotheses and used formulae.

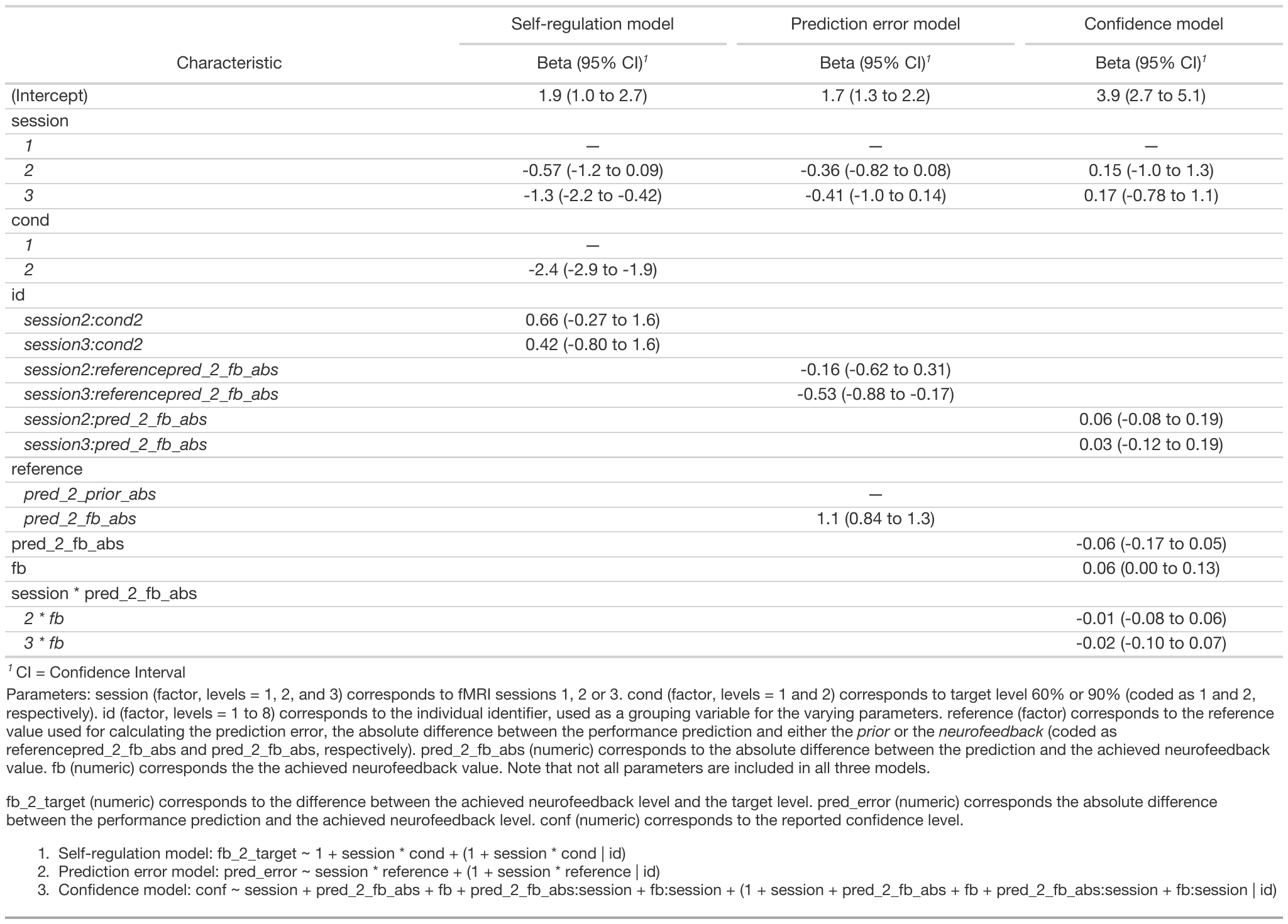

**Table S3**. Summaries of tested models.

Whole-brain fMRI analysis: resulting clusters

| Contrast | x | y | z | Size | Area |
| --- | --- | --- | --- | --- | --- |
| Regulation ((Level 60% + Level 90%) > rest) | 39 | -36 | 40 | 1747 | Right: supramarginal gyrus |
|  | 29 | 1 | 13 | 10.299 | Right: thalamus, basal ganglia, insula, inferior frontal gyrus, middle frontal gyrus |
|  | -14 | -6 | 38 | 32.938 | Bilateral: premotor areas, SMA |
|  | 38 | 31 | 33 | 909 | Right: dorsolateral prefrontal cortex |
|  | 26 | -54 | -22 | 2.744 | Right: cerebellum |
|  | 21 | -66 | 42 | 2.354 | Right: superior parietal lobe, precuneus |
|  | -29 | -56 | 42 | 6.205 | Left: superior parietal lobe, precuneus |
|  | -22 | -56 | -24 | 936 | Left: cerebellum |
|  | -41 | 29 | 25 | 561 | Left: dorsolateral prefrontal cortex |
|  | -56 | -30 | 27 | 709 | Left: supramarginal gyrus |
| Performance prediction >  Performance prediction catch | 0 | 38 | 20 | 195.723 | Bilateral: thalamus, subthalamic nucleus, substania nigra, red nucleus, putamen, medial and lateral globus pallidus, dentate, caudate, cerebellum, fusiform gyrus, parahippocampal gyrus, cingulate gyrus, cuneus, precuneus, middle and inferior occipital gyrus, superior and inferior parietal lobule, postcentral gyrus, paracentral lobe, precentral gyrus, inferior, superior and middle frontal gyrus, supramarginal gyrus, lingual gyrus, insula. |
|  | 39 | 31 | 31 | 4.659 | Right: inferior and middle frontal gyrus, precentral frontal gyrus |
|  | 11 | -38 | 44 | 216 | Right precuneus and right cingulate gyrus |
|  | 1 | 38 | 30 | 2.197 | Bilateral: cingulate gyrus and posterior cingulate |
|  | -34 | 38 | 30 | 1.212 | Bilateral: Cerebellum (dentate, cerebellar lingual, culmen, declive, fastigium, nodule, pyramis, tuber, uvula), brain stem (pons, substrantia nigra, red nucleus, geniculum body, subthalamic nucleus), angular gyrus, caudate, cingulate gyrus, claustrum, cuneus, precuneus, corpus callosum, fusiform gyrus, inferior, medial, and superior frontal gyrus, inferior and middle occipital gyrus, inferior and superior parietal lobule, insula, lentiform nucleus (globus pallidus, putamen), lingual gyrus, middle temporal gyrus, paracentral lobule, parahippocampal gyrus, precentral gyrus, postcentral gyrus, posterior cingulate, supramarginal gyrus, thalamus (mammillary body, ventral lateral and ventral posterior medial, medial dorsal, ventral anterior, anterior nucleus, pulvinar, medial geniculum body);  Right superior occipital gyrus; left superior temporal gyrus |
| Confidence > Confidence catch | 28 | -73 | 7 | 19.005 | Right: Cerebellum (declive, culmen), cuneus, fusiform gyrus, inferior, middle, and superior occipital gyrus, lingual gyrus, middle temporal gyrus, precuneus, superior parietal lobule, supramarginal gyrus |
|  | 41 | -2 | 35 | 396 | Right: middle frontal gyrus, precentral gyrus |
|  | 42 | 11 | 13 | 236 | Right: insula, precentral gyrus |
|  | 32 | 22 | 9 | 264 | Right: insula, inferior frontal gyrus |
|  | 25 | -8 | 52 | 695 | Right: middle frontal gyrus, precentral gyrus |
|  | 20 | -26 | 3 | 1.421 | Right: thalamus (pulvinar), parahippocampal gyrus |
|  | 10 | 3 | 5 | 108 | Right caudate |
|  | 1 | -22 | -8 | 2.458 | Bilateral midbrain (substantia nigra, red nucleus, subthalamic nucleus, medial dorsal nucleus) |
|  | 3 | -63 | -23 | 522 | Cerebellum |
|  | -28 | -77 | -5 | 13.146 | Left: cerebellum (culmen, declive, tuber, uvula), cuneus, fusiform gyrus, inferior and middle occipital gyrus, lingual gyrus, middle temporal gyrus, precuneus |
|  | -14 | -22 | 11 | 111 | Left thalamus (pulvinar, medial dorsal and lateral posterior nuclei) |
|  | -21 | -27 | -1 | 566 | Left: lateral geniculum body, parahippocampal gyrus, thalamus (pulvinar) |
|  | -26 | -60 | 42 | 1.370 | Left: inferior parietal and superior lobule, precuneus |
|  | -27 | -12 | 46 | 183 | Left: middle frontal gyrus, precentral gyrus |

**Table S4**. fMRI results for contrasts Regulation (60% + 90%) > Baseline, Performance prediction > Performance prediction catch, and Confidence > Confidence catch. For each significant cluster, the coordinates of the center of gravity are reported, as well as the size of it, and the regions included in the cluster.

Parametric modulation

To further explore the data, parametric effects of our two main self-evaluation conditions, the performance prediction and reports of confidence, were also investigated. The parametric weights were inserted on the corresponding scale from 0 to 12 for performance prediction and 0 to 10 for confidence (0 being 50% and 10 being 100% confidence). These values were normalized by subtracting the corresponding session’s average report. A fixed-effects general linear model including 8 conditions (same as for self-evaluation conditions (see main text), but with the addition of either the parametric condition of performance prediction or confidence) was calculated for each participant separately. Z-transformed motion predictors were again included as confounds. An FDR-corrected t-map (*q* = 0.05) of the conjunction contrast of the main and parametric predictor (i.e., performance prediction or confidence) was created for each participant and a probability map was generated for all 8 participants for each of the two conditions.

Since the resulting two probability maps did not result in any surviving clusters when including at least 50% of the participants, we performed some further exploratory analyses. We first wanted to see if accuracy of prediction (prediction – neurofeedback) or the difference between the target level and prediction resulted in any parametric effects. Each difference was included as an additional regressor in a separate fixed-effects general linear model using the timepoints of the performance prediction timeframe, following the same procedure as described above. These two probability maps also did not reveal any surviving clusters.

Finally, as none of the previous analyses included surviving clusters, we again generated all four probability maps described above, but this time using a mask to decrease the number of multiple comparisons. The mask was based on a meta-analysis of different metacognitive processes reported in 47 studies (Vaccaro & Fleming, 2018) and consisted of 8 spheres, each with the center in the peak coordinate reported in the meta-analysis (see figure S4 and table S3) and with a liberal radius of 10 voxels (~33.500 mm3).

*Results.* In order to see if the performance predictions or confidence reports would be reflected in the strength of the brain activation, we also performed GLM using parametric weights for each individual subject. Four probabilistic maps, each including one additional parametric condition, were created with two different coverages: full-brain coverage or with a mask. The four additional conditions included either performance prediction, confidence reports, accuracy of prediction (i.e., prediction – neurofeedback), or error prediction (i.e., performance prediction – target level). None of the eight resulting probabilistic maps showed any parametric effects.

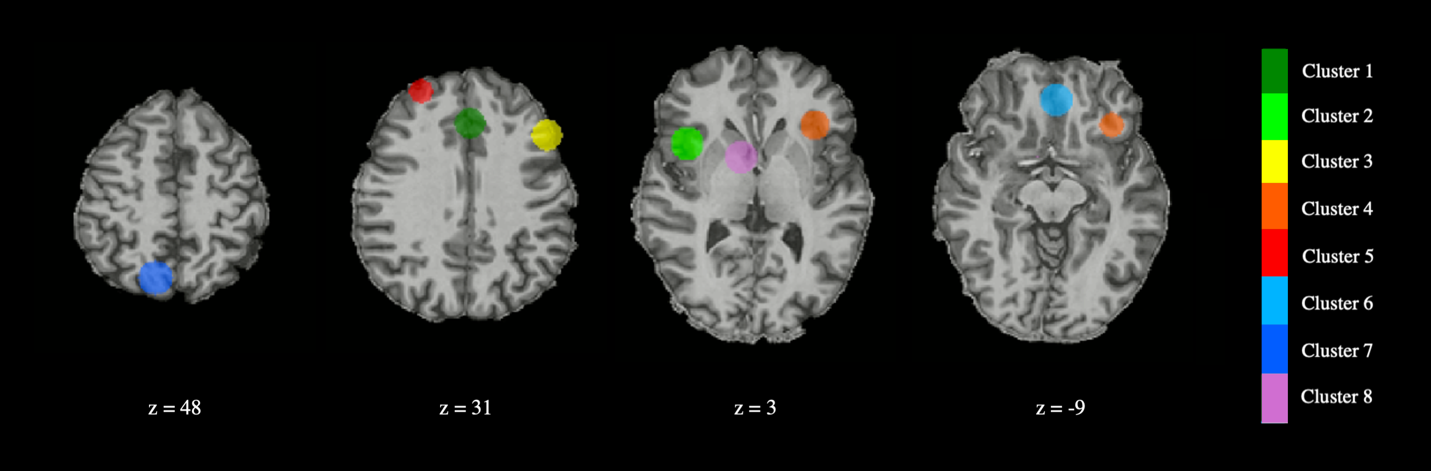

**Figure S4**. Regions included in the mask. Each of the spheres correspond to one cluster in Vaccaro & Fleming (2018); its center in positioned in the peak coordinate (see Table S3).

| **Cluster** | **Central coordinate (TAL)** | | | **Region** |
| --- | --- | --- | --- | --- |
|  | **x** | **y** | **z** |  |
| 1 | -2 | 30 | 35 | L/R posterior medial frontal cortex |
| 2 | 41 | 13 | 4 | R insula/inferior frontal gyrus |
| 3 | -48 | 23 | 27 | L dorsolateral prefrontal cortex |
| 4 | -35 | 24 | -2 | L insula/inferior frontal gyrus |
| 5 | 27 | 49 | 24 | R anterior dorsolateral prefrontal cortex |
| 6 | -2 | 39 | -12 | L/R ventromedial prefrontal cortex |
| 7 | 12 | -62 | 48 | R dorsal precuneus |
| 8 | 9 | 5 | 1 | R ventral striatum |

**Table S5.** The regions and their central coordinate used to create a mask for GLM modelling. The central voxel coordinates are based on the peak coordinates reported in Vaccaro & Fleming (2018). Note that the region naming corresponds to the one in the original publication but might include neighboring regions in the present report due to the spherical nature of clusters. Note also that the original peak coordinates were reported in MNI space. Transformations from MNI to Tal coordinates needed for this study were performed using an online application from BioImage Suite, found here: https://bioimagesuiteweb.github.io/webapp/mni2tal.html. L = left; R = right.
